# Therapeutic *Salmonella* induces long-term protective trained immunity in NK cells against cancer metastasis

**DOI:** 10.1101/2025.08.29.673007

**Authors:** Li Rong, Jingchu Hu, Wei Jin, Renhao Li, Yangfan Wu, Nan Zhou, Zhongjun Zhou, Hosen Shakhawat, Xiaolei Wang, Shuo Han, Huarui Gong, Xuansheng Lin, Wenjun Li, Yige He, Sukbum Kim, Xiaoyu Zhao, Qiubin Lin, Jian-Dong Huang

**Affiliations:** State Key Laboratory of Quantitative Synthetic Biology, Shenzhen Institute of Synthetic Biology, Shenzhen Institutes of Advanced Technology, Chinese Academy of Sciences, Shenzhen 518055, China; School of Biomedical Sciences, Li Ka Shing Faculty of Medicine, The University of Hong Kong, Pokfulam, Hong Kong SAR, China; Department of Medicine, School of Clinical Medicine, Li Ka Shing Faculty of Medicine, The University of Hong Kong; Pokfulam, Hong Kong SAR, P.R. China; ZJU-Hangzhou Global Scientific and Technological Innovation Center, Zhejiang University, Hangzhou 311215, China; Shenzhen Key Laboratory for Cancer Metastasis and Personalized Therapy, Department of Clinical Oncology, The University of Hong Kong-Shenzhen Hospital, Shenzhen 518053, China; Guangdong-Hong Kong Joint Laboratory for RNA Medicine, Sun Yat-Sen University, Guangzhou 510120, China; Materials Innovation Institute for Life Sciences and Energy (MILES), HKU-SIRI, Shenzhen, P.R. China

**Keywords:** trained immunity, NK cells, metastasis, *Salmonella*

## Abstract

Trained immunity, a form of innate immune memory initially recognized in pathogen defense, has recently been reported to aid in combating cancer. Although trained immunity has been well-characterized in myeloid lineage cells, its presence in innate lymphoid cells, particularly natural killer (NK) cells, remains largely unexplored. Here, we report that a single dose of *Salmonella* YB1, an attenuated candidate anticancer vaccine strain, provides long-lasting protection against cancer metastasis in mice by inducing trained natural killer (NK) cells. Functional assays coupled with multiomics analysis revealed that *Salmonella*-trained NK (stNK) cells establish an enduring reprogrammed epigenome characterized by enhanced pro-survival signaling and immune effector functions, resulting in more potent IFN-γ release and cytotoxicity upon secondary stimulation. We further revealed that IL-12 and IL-18 are essential but insufficient for this NK cell training effect. Crucially, stNK cells far outperform the common immune checkpoint therapies, including PD-1 and TIGIT blockade, in suppressing metastasis, underscoring the unique immunological mechanisms in combating metastasis. These findings uncover a novel long-term anti-metastatic function of trained immunity in NK cells, highlighting its potential as an effective antitumor strategy.

## Introduction

Cancer immunotherapy has revolutionized the cancer treatment field, offering various agents to mobilize adaptive antitumor immunity, including bispecific antibodies, immune checkpoint inhibitors, and chimeric antigen receptors. Despite their success, these therapies only benefit a minority of patients, emphasizing the need to develop next-generation immunotherapies. Given the crucial role of innate immune responses in immunity, leveraging innate immune cells offers great potential for durable and multifaceted tumor control[1].

Trained immunity, a long-term functional reprogramming of innate immune cells, is characterized by enhanced responses of innate immune cells to a secondary challenge after returning to a non-activated state, which results from enduring epigenetic reprogramming[2]. Trained immunity was initially and has been extensively studied in the context of host defense against infectious diseases in humans and experimental animals. In these studies, myeloid lineage cells, including monocytes, macrophages, and their precursor cells, were conditioned by specific stimuli such as the Bacillus Calmette–Guérin (BCG) vaccine, β-glucan, or LPS, to boost protection against microbial pathogens[2–6]. Recent studies have reported the involvement of trained immunity in anti-tumor immune response, primarily focusing on myeloid cells and their direct or indirect effects on tumor cell elimination[7–10]. However, it remains poorly explored whether known or undiscovered stimuli can induce trained immunity in innate lymphoid cells (ILCs) and, if so, whether such reprogramming confers anti-tumor effects in these cells[11].

Derived from the bone marrow, NK cells are the sole cytotoxic subset among ILCs and are abundantly present in blood and lymphoid tissues, with a rapid turnover of approximately two weeks[12]. NK cells possess natural anti-tumor capabilities, which include direct cytotoxic attack on tumor cells and modulation of the tumor microenvironment via secretion of anti-tumoral cytokines such as IFN-γ and TNF-α. These multifaceted capabilities make NK cells highly promising candidates for effective cancer immunotherapy[13]. Emerging studies indicate that NK cells can develop antigen-specific immunological memory in response to some stimuli, with mouse cytomegalovirus (MCMV) and human cytomegalovirus (HCMV) infection being the most well-studied examples. Mouse NK cells expressing the Ly49H receptor and human NK cells expressing the NKG2C receptor possess antigen specificity for MCMV-encoded glycoprotein m157 and HCMV-encoded UL40 peptides, respectively. Once engaged with the stimuli, these NK cells sequentially undergo clonal proliferation and contraction, and finally establish long-term persistence akin to memory T cells, with the prerequisites for pro-inflammatory cytokines, such as IL-12 and IL-18. Because such NK cell responses more closely resemble the antigen-specific T cell response instead of the myeloid cell trained immunity, these NK cells are referred to as memory NK cells. In contrast, overnight priming of NK cells *in vitro* solely with a combination of IL-12, IL-15, and IL-18 endows memory-like features characterized by increased IFN-γ production, but no changes in cytotoxicity upon non-antigen-specific re-stimulation. These NK cells, termed cytokine-induced memory-like (CIML) NK cells, more closely align with the concept of trained immunity[2, 14, 15]. Despite these findings, it remains to be explored whether trained immunity exists in NK cells *in vivo* and, if so, whether it can be harnessed to develop a long-lasting antitumor therapy.

*Salmonella* YB1, engineered from the *S. Typhimurium* SL7207, exhibits attenuated virulence while retaining anti-tumor efficacy[16]. We previously demonstrated that *Salmonella* YB1 infection induces acute inflammation and effectively suppresses cancer metastasis within a short period (two weeks) post-infection by activating NK cells [17, 18]. However, it remains unclear whether the *Salmonella* YB1-treated animals develop long-lasting protective immunity beyond the general NK cell turnover time (around 2 weeks) and, if so, what the underlying cellular and molecular mechanisms are.

Here, we report that a single dose of *Salmonella* YB1 confers long-term protection against metastasis across diverse mouse genetic backgrounds and cancer types by inducing persistent trained immunity in peripheral NK cells. Mechanistically, *Salmonella*-trained NK (stNK) cells continue to exhibit enhanced IFN-γ release and cytotoxicity upon secondary stimulation after the resolution of anti-infectious inflammation, which correlates with and is most likely attributed to the sustained epigenetic reprogramming in these NK cells. Further investigation indicates that IL-12 and IL-18 are essential but insufficient for this NK cell training effect. Strikingly, stNK cells outperform common immune checkpoint therapies (e.g., PD-1 blockade [19], TIGIT blockade[20]) in suppressing metastasis. These findings reveal a broad-spectrum, durable anti-metastatic function of stNK cells, reinforcing the concept of trained immunity by expanding the range of stimuli, the types of immune cells being trained, and the functional scenarios, and highlighting the potential of harnessing trained NK cells to combat deadly cancer metastasis in future clinical therapies.

## Results

### Attenuated *Salmonella* induces long-lasting anti-metastatic immunity

To determine whether the *Salmonella* YB1-elicited anti-metastatic effect persists beyond acute inflammation phase, we first quantified the duration of the inflammation. Wild-type (WT) BALB/c and C57BL/6J mice were intravenously (*i.v.*) injected with *Salmonella* YB1 or phosphate-buffered saline (PBS), and their body weight changes were monitored. The results revealed that the initial body weight loss caused by YB1 treatment was restored within two weeks (**Figure S1a, b**). These findings, along with the rapid clearance of YB1 within a week post-infection[18] and the gradual normalization of elevated pro-inflammatory cytokines (such as IL-18, IL-6, and TNF-α) to baseline levels within 18 days post-infection **(Figure S1c)**, suggest that YB1-induced acute inflammation persists for less than one month.

To examine the long-term effects of *Salmonella* YB1 treatment, BALB/c mice were *i.v.* injected with murine mammary carcinoma 4T1 cells during the acute infection phase (on day 6 post-infection) or after the inflammation resolution phase (on days 30, 60, 90, and 120 post-infection) to develop lung metastasis (**Figure 1a**). The results revealed that YB1-treated mice not only exhibited potent anti-metastatic ability during the acute infection phase, but also retained this protection for several months afterward, suggesting a durable anti-metastatic immunity (**Figure 1b-d**). To save time, we chose day 30 post-infection as a representative long-term time point for further mechanistic investigations. Luciferase-labeled 4T1 cells (4T1-Luci) were *i.v.* injected into BALB/c mice pretreated with YB1 or PBS 30 days in advance and the *in vivo* presence of 4T1-Luci cells was subsequently monitored using luciferase live imaging. 24 hours after the injection of the 4T1-Luci cells, a significant accumulation of luciferase signal was observed in the lungs of PBS-treated mice, whereas the signal was notably diminished in YB1-treated mice **(Figure 1e, f; Figure S1d**). This observation demonstrates that the long-term anti-metastatic immunity induced by *Salmonella* YB1 treatment interferes with the colonization process of the metastatic cascade, particularly during the early cancer cell survival phase.

**Figure 1.**
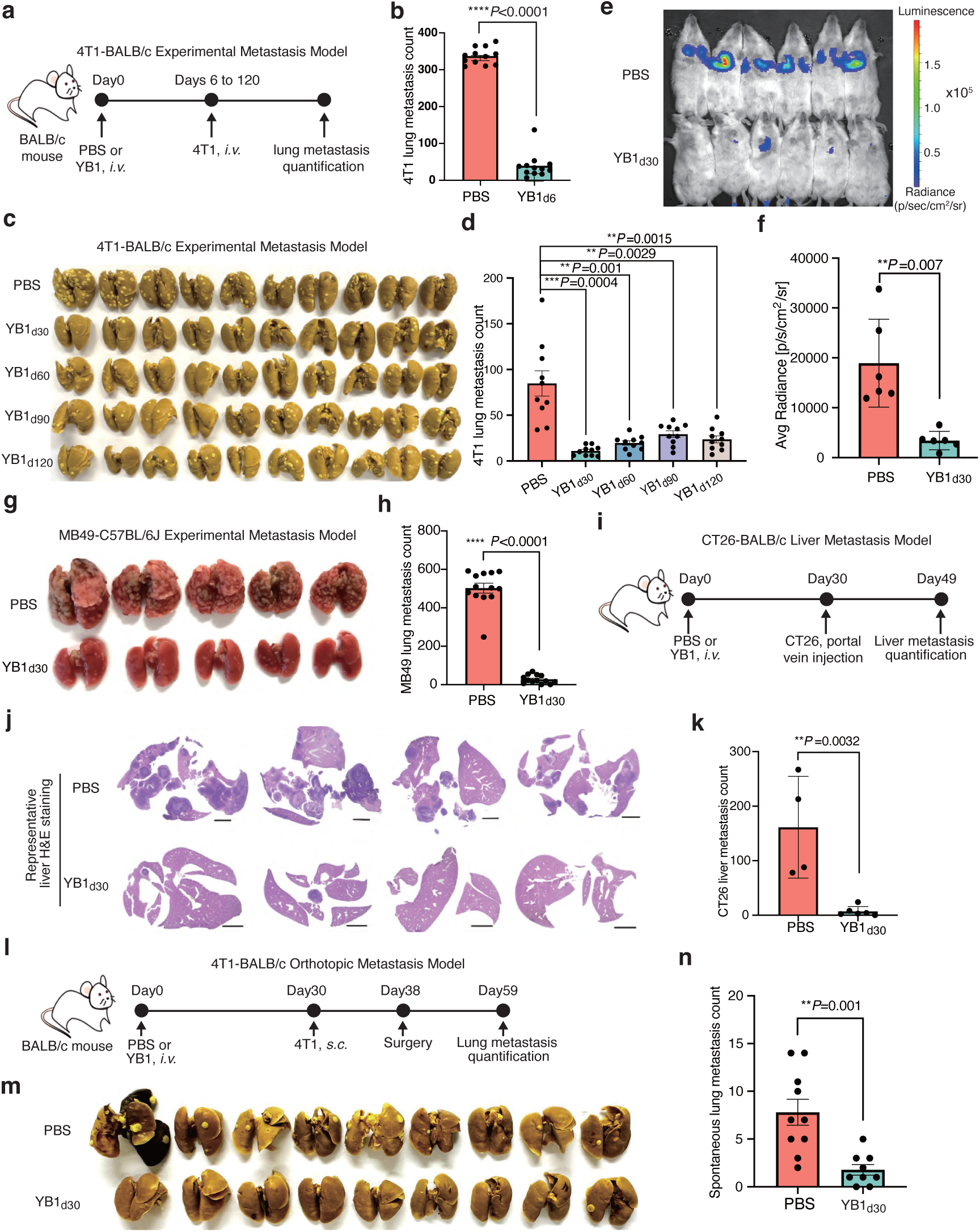
Attenuated *Salmonella* induces long-lasting anti-metastatic immunity. The subscript number of YB1 indicates the number of days post-infection with *Salmonella* YB1 at the time of tumor cell inoculation for Figure 1a-n. **a)** A schematic representation of *Salmonella* YB1 treatment and 4T1-BALB/c experimental metastasis model. 4T1 cancer cells were *i.v.* inoculated at different time points, ranging from 6 days to 120 days post-YB1 infection, to develop lung metastasis burden. PBS treatment serves as a control. **b)** Quantification of 4T1 lung metastasis (n=12 per group) for the 4T1 cancer cells inoculation occurred on day 6 post-YB1 or -PBS treatment. **c-d)** Images of Bouin-fixed lungs and quantification of 4T1 lung metastases (n = 10 per group) to validate the long-term anti-metastatic effects of YB1 treatment, as indicated in Figure 1a. **e-f)** Tracking and quantification (n=6 per group) of 4T1-Luci cells *in vivo* using luciferase live imaging. **g-h)** In the MB49-C57BL/6J experimental metastasis model, MB49 bladder cancer cells were *i.v.* injected into C57BL/6J mice pretreated with either PBS or YB1 30 days prior to evaluate the anti-metastatic efficacy. The representative image of 4% PFA-fixed lungs and quantification of MB49 lung metastatic nodules (n=13 per group) are shown here. **i)** A schematic representation of CT26-BALB/c experimental liver metastasis model. **j-k)** H&E staining of whole liver tissue (scale bar, 2mm) and quantification of liver metastatic nodules (n = 4 for PBS, n=6 for YB1_d30_) as indicated in Figure 1I. Condensed tumor nodules were stained with H&E from PBS samples. **l)** A schematic representation of 4T1-BALB/c orthotopic metastasis models. The post-surgery lung metastasis was analyzed as demonstrated. **m-n)** Images of Bouin-fixed lungs and quantification of 4T1 lung metastases (n =10 for PBS, n=9 for YB1_d30_) as indicated in Figure 1l. **Figure 1b, d, h, and n** are displayed as a combined result of two independent experiments and presented as mean values ± SEM. **Figure 1f, k** are displayed as one representative experiment of two independent experiments and presented as mean values ± SD. All *P* values are yielded by two-tailed unpaired t-tests and are shown in the relevant figure.

YB1 treatment also elicits long-lasting anti-metastatic immunity in the experimental metastasis model established with MB49 bladder cancer cells and C57BL/6 mice, confirming YB1’s long-term effects are independent of the cancer types or the host genetic backgrounds (**Figure 1g, h**; **Figure S1e, f**). We further confirmed that YB1-induced long-term anti-metastatic immunity also applies to a liver metastasis model established in BALB/c mice through portal vein injection of CT26 colon cancer cells (**Figure 1i-k; Figure S1g**). To simulate the clinical scenario of post-surgical metastasis in patients with newly diagnosed breast cancer, we established an orthotopic model of breast cancer metastasis (**Figure 1l**). In this model, mice were initially treated with YB1, followed by inoculation of 4T1 cancer cells into the mammary fat pad 30 days later to develop orthotopic breast tumors. Subsequently, visible orthotopic breast tumors (approximately 50 mm^3^) were surgically removed 8 days later, and the lung metastasis post-surgery was quantitatively analyzed. Under such condition, YB1-induced long-term protective immunity effectively prevented metastasis following surgery (**Figure 1l-n**). These data collectively indicate that *Salmonella* YB1 treatment induces durable systemic anti-metastatic immunity in various syngeneic mouse metastasis models, even after the resolution of anti-infectious inflammation.

### NK cells mediate the long-lasting anti-metastatic immunity conferred by the attenuated *Salmonella* treatment

The long-term protective effects of the immune system following the return from acute inflammation phase to homeostasis, known as immunological memory, have long been attributed to the adaptive immune responses typically featured by the participation of T and B lymphocytes. We observed that YB1 treatment still significantly reduced the MB49 lung metastatic burden in B6 *Rag1* knockout mice — which lack mature T and B cells[21] — even at 30 days post-infection (**Figure 2a, b**), indicating that adaptive immunity is dispensable for the establishment of YB1-induced long-lasting anti-metastatic immunity. Compared to the T and B cells totally depleted *Rag1* knockout mice, NSG mice exhibit broader cellular deficiencies, particularly in NK cells[22], thus were selected for NK cell role analysis. Due to genetic background differences between *Rag1* knockout and NSG mice, here we employed the athymic nude mice as the control as they have the same genetic background as NSG mice and lack T cells but have intact NK cells [23, 24]. Our results showed that YB1 treatment significantly reduced the lung metastatic burden in nude mice, whereas there was negligible difference in the lung metastatic burden between the YB1-treated and PBS-treated NSG mice (**Figure 2c-e**). Furthermore, both the YB1-treated and PBS-treated NSG mice exhibited a lung metastatic burden (**Figure 2d**) approximately 40 times higher than that of the PBS-treated nude mice (**Figure 2e**). These data suggest that the absence of NK cells greatly diminishes the host’s anti-metastatic ability, and the long-lasting anti-metastatic capability induced by YB1 most likely mainly rely on NK cells, but not T and B cells.

**Figure 2.**
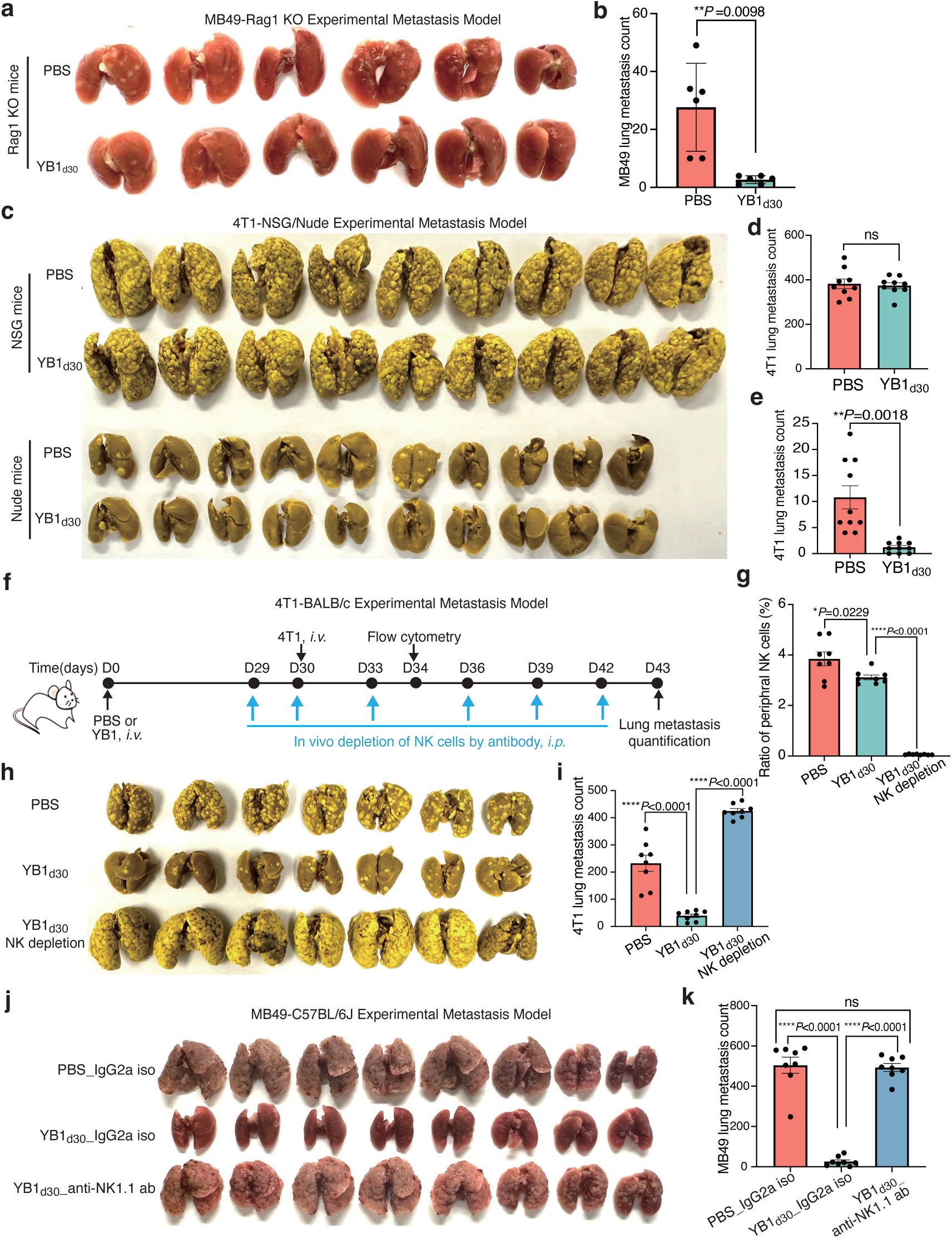
NK cells mediate the long-lasting anti-metastatic immunity conferred by attenuated *Salmonella* treatment. **a-b)** MB49 bladder cancer cells were *i.v.* injected into *Rag1* knockout mice pretreated with either PBS or YB1 30 days prior to evaluate the anti-metastatic efficacy. Images of 4% PFA-fixed lungs and quantification of MB49 lung metastatic nodules (n=6 per group, presented as mean values ± SD) are shown here. **c-e)** The same dosage of 4T1 cancer cells were *i.v.* injected into each NSG mouse and Nude mouse pretreated with either PBS or YB1 30 days prior to evaluate the anti-metastatic efficacy. Images of Bouin-fixed lungs (**c**) and the quantification of 4T1 lung metastases (n=9 per group in NSG mice, n=10 per group in Nude mice) in NSG mice (**d**) and Nude mice (**e**) are presented. **f)** A schematic representation of continuous NK cell depletion schedule using Anti-asialo GM1 antibody in BALB/c mice pretreated with YB1 30 days prior. **g)** The peripheral ratio of NK cells was validated by flow cytometry (n= 8 per group) 4 days after 4T1 cancer cell inoculation, as depicted in Figure 2f. **h-i)** Images and quantification (n=8 per group) of 4T1 lung metastases, as depicted in **Figure 2f**. **j-k)** C57BL/6J mice were treated with PBS, YB1, or YB1 plus NK cell depletion antibody (Anti-NK1.1 antibody), respectively. MB49 cancer cells were *i.v.* injected 30 days post-treatment with either YB1 or PBS, as depicted in **Figure S2c.** Images and quantification of MB49 lung metastases (n = 8 per group) across three groups are presented here. All quantified data are shown as combined results of 2 independent experiments and presented as mean values ± SEM unless indicated. All *P* values are yielded by two-tailed unpaired t-tests and are shown in the relevant figure.

To further ascertain the role of NK cells, we depleted the NK cells in BALB/c mice by injecting the anti-asialo-GM1 antibody as previously described with a minor modification[25]. Detailly, NK cell depletion was initiated on day 29 after YB1 treatment and continued until the experimental endpoint, wherein 4T1 tumor cells were inoculated through the tail vein on day 30 (**Figure 2f**). The efficiency of NK depletion *in vivo* was confirmed by the absence of peripheral CD3− NKp46+ CD49b+ cells assayed by flow cytometry (**Figure 2g**). The gating strategies for identifying NK cells are presented in **Figure S2a**. We observed that the depletion of NK cells abolished the YB1-induced long-lasting anti-metastatic immunity and even led to more metastatic nodules in the lungs compared to the PBS group (**Figure 2h, i; Figure S2b**). To double confirm it, we also repeated the NK cell depletion experiment with the anti-NK1.1 antibody in C57BL/6J mice, which showed similar results (**Figure 2j, k**; **Figure S2c-f**). Collectively, these data demonstrated that the long-lasting anti-metastatic effects induced by *Salmonella* YB1 treatment are mainly mediated by NK cells.

### Attenuated *Salmonella*-trained NK cells undergo robust population remodeling and exhibit enhanced effector responses upon secondary stimulation

Given that NK cells trained by YB1 treatment not only suppress metastasis during the acute infection phase but also maintain this anti-metastatic effect long-term after the resolution of anti-infectious inflammation, it is imperative to investigate their quantitative and functional changes over time.

During the acute phase of YB1 infection, the ratio of peripheral NK cell declined within 3 hours, gradually rebounded to baseline level (PBS-treated mice) by 2 days, and surpassed baseline thereafter (**Figure 3a**). Consistent with the NK cell population dynamics, *in vivo* bioluminescence imaging tracking 4T1-Luci tumor cells indicated that it was only starting from day 2 post-infection that YB1-treated mice exhibited a stronger potential to eliminate metastatic tumor cells (**Figure 3b, c**). These findings demonstrate that during the acute infection phase, the NK cell population undergoes a transient decline followed by replenishment within 2 days, with the newly replenished NK cells exhibiting enhanced anti-tumor activity.

**Figure 3.**
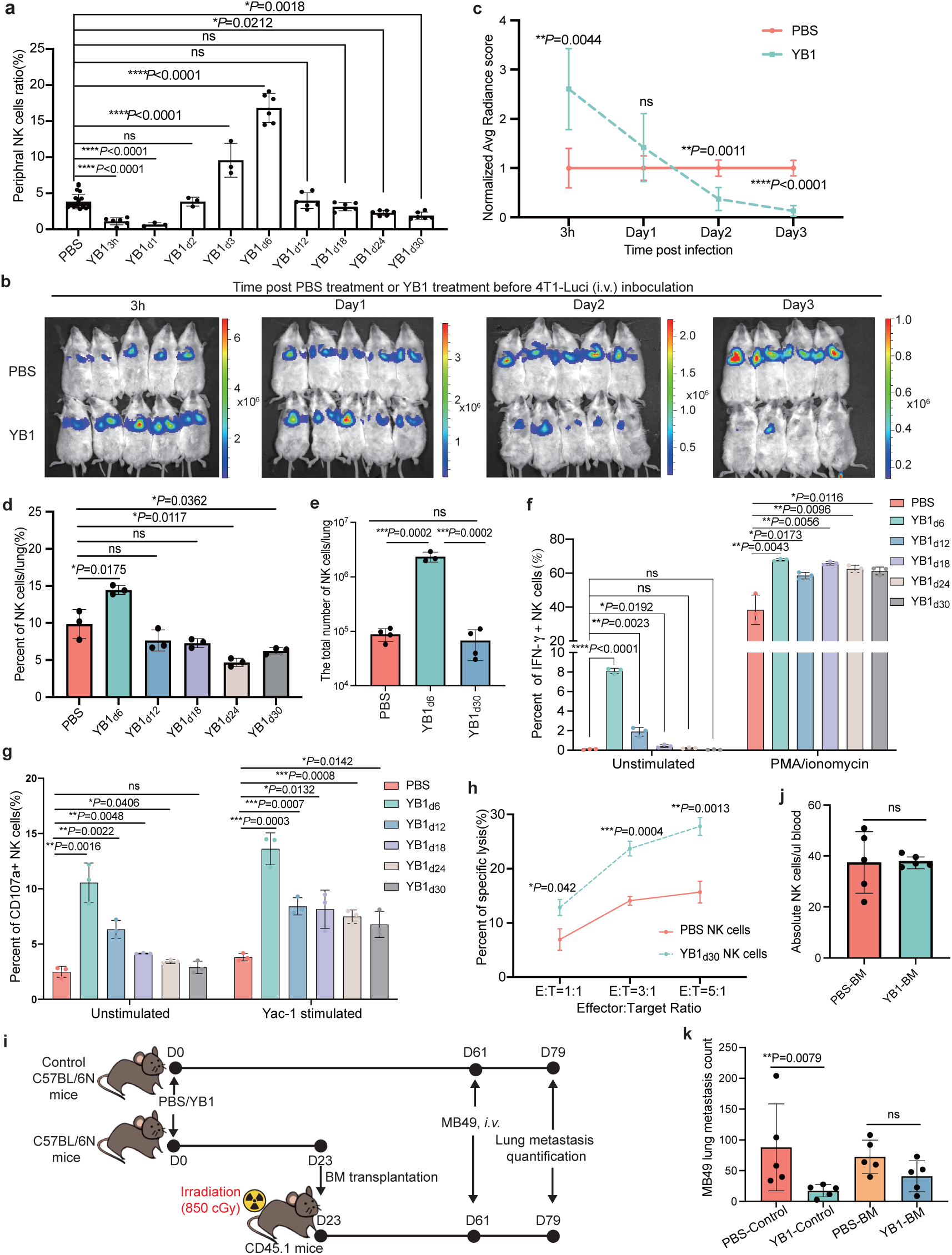
Attenuated *Salmonella*-trained NK cells undergo robust population remodeling and exhibit enhanced effector responses upon secondary stimulation. **a)** The percentage of peripheral NK cells at different time points post YB1 treatment in BALB/c mice, as determined by flow cytometry (n=22 for PBS group, n=6 for YB1_3h_/YB1_d6_/YB1_12_/YB1_18_/YB1_24_ /YB1_30,_ n=3 for YB1_d1_/YB1_d2_/YB1_d3_, combined results of 4 independent experiments, data are presented as mean values ± SD). PBS serves as a control. **b-c)** Tracking and normalized quantification (n≥4 per group) of 4T1-Luci cells *in vivo* by luciferase live imaging 3 hours after *i.v.* injection of these cells into BALB/c mice. The mice were pretreated with either PBS or YB1 at different time points (3 hours, 1 day, 2 days, and 3 days) prior to the inoculation of 4T1-Luci cancer cells. **d)** The percentage of NK cells to all immune cells in the lung at different time points post YB1 treatment in BALB/c mice, as determined by flow cytometry (n=3 per group). PBS serves as a control. **e)** The absolute total number of NK cells per lung across samples (n≥3 per group). Total immune cell numbers were measured by trypan blue exclusion and then multiplied by the percentage of NK cells determined by FACS analysis to give the absolute number of NK cells for each lung. **f)** Flow cytometric analysis was performed to assess the *ex vivo* production of IFN-γ in lung-infiltrating NK cells, both in the absence and presence of PMA/Ionomycin stimulation for 5 hours *ex vivo* (n= 3 per group). **g)** Flow cytometric analysis was conducted to evaluate CD107a expression on lung-infiltrating NK cells both in the absence and presence of co-culture with YAC-1 cells for 5 hours *ex vivo* (n=3 per group). **h)** Cytotoxicity of lung-infiltrating NK cells against YAC-1 target cells at the indicated NK cells: YAC-1 cells (E: T, Effector: Target) ratios (n=3 biological replicates). **i)** A schematic representation of bone marrow transplantation. **j)** Measurement of donor-derived NK cell concentration (cells/μL of blood) following a 5-week bone marrow reconstitution period, as depicted in Figure 3i. **k)** Quantification (n=5 per group) of MB49 lung metastasis, as depicted in Figure 3i. One representative experiment of two independent experiments is displayed and presented as mean values ± SD for Figure 3c-k. All *P* values are yielded by two-tailed unpaired t-tests and are shown in the relevant figure.

The inflammation induced by YB1 was estimated to subside quickly (**Figure S1a-c**). Consistently, the increase of peripheral and lung-infiltrating NK cells following YB1 treatment gradually returned to the baseline level by day 12 and continued to decline thereafter (**Figure 3a, d**). Although the ratio of NK cells in YB1-treated mice was lower than the baseline level on day 30 post-infection, there was no significant change in the absolute number of lung-infiltrating NK cells between PBS- or YB1-treated mice (**Figure 3d, e**). Moreover, proliferating NK cells labeled with Edu during the early acute infection phase could still be detected in YB1-treated mice 30 days post-infection (**Figure S3b, c**). These findings collectively suggest that following *Salmonella* YB1 treatment, NK cells sequentially undergo the proliferation, contraction, and long-term persistence phases, exhibiting similar processes as the formation of memory T cells[26].

Despite the restoration of NK cell number to the baseline level 30 days post-YB1 treatment, the YB1-treated mice still exhibited stronger metastasis suppression than PBS-treated mice (**Figure 2**), raising the question of whether these YB1-trained NK cells remained in a continuously activated state. We isolated lung-infiltrating NK cells at various time points after YB1 treatment and evaluated their *ex vivo* production of IFN-γ and the cell-surface level of CD107a as a surrogate for degranulation (**Figure f, g**; **Figure S3d, e**). NK cells isolated from YB1-treated mice during the acute infection phase (day 6) indeed exhibited hyperactivity, as evidenced by spontaneous secretion of IFN-γ and elevated level of CD107a in the absence of any stimulus during the *ex vivo* analysis. However, this hyperactive state completely subsided by day 24 post-YB1 infection, indicating that these NK cells had restored homeostasis. Interestingly, we observed that the *Salmonella*-trained NK (stNK) cells that had already returned to homeostasis were able to produce more IFN-γ upon PMA/Ionomycin stimulation and expressed higher levels of CD107a in response to co-culturing with Yac-1 cancer cells. In line with the elevated CD107a levels, the stNK cells that had restored homeostasis 30 days post-infection also exhibited higher cytotoxicity against Yac-1 cancer cells upon co-culturing (**Figure 3h; Figure S3f**). These observations reveal enhanced responses of stNK cells compared to control NK cells upon secondary stimulation after the inflammation resolution, which highly suggests that *Salmonella* YB1 treatment induces long-lasting trained immunity in NK cells.

To investigate the role of central trained immunity, manifested as trained bone marrow-resident progenitors continuously generating functionally committed anti-metastatic NK cells, in YB1-induced long-term anti-metastatic protection, we conducted bone marrow transplantation experiments (**Figure 3i-k; Figure S4a-e**). Bone marrow cells extracted from PBS- or YB1-treated C57BL/6N mice were transplanted into naïve irradiated syngeneic CD45.1 mice, which achieved successful reconstitution 5 weeks later (**Figure 3i, j; Figure S4a-c**). Then reconstituted recipient mice and control C57BL/6N mice were *i.v.* injected with MB49 bladder cancer cells (**Figure 3i**). In contrast to control C57BL/6N mice, there was no significant difference in MB49 tumor burden in the lungs of mice receiving bone marrow from PBS-treated versus YB1-treated mice (**Figure 3k; Figure S4d, e**). These findings imply that central trained immunity is not involved in the long-term protection against metastasis following *Salmonella* YB1 treatment.

Collectively, these results demonstrate that *Salmonella* YB1 treatment induces NK cell proliferation and activation during acute infection, leading to robust suppression of metastasis in this phase. Subsequently, as inflammation resolves, the NK cell population contracts to baseline levels while concurrently exiting the activated state. Thereafter, the stabilized NK cells persist long-term in tissues and retain enhanced responsiveness to secondary stimulation.

### Attenuated *Salmonella* induces long-term epigenetic reprogramming characteristic of trained immunity in NK cells

Given that NK cells cannot undergo somatic rearrangement, such as V(D)J recombination seen in T and B cells, their enhanced functions are likely driven by sustained epigenome remodeling. To investigate the epigenome landscape of the stNK cells, lung-infiltrating NK cells from YB1-treated mice during the acute infection phase (6 days post-infection) and after the resolution of the anti-infectious inflammation (30 days post-infection) were isolated and subjected to chromatin accessibility analysis using ATAC-seq (assay for transposase-accessible chromatin with high throughput sequencing). Compared with the NK cells from the PBS-treated group, on day 6 post-infection, the stNK cells exhibited significantly increased chromatin accessibility at 16,806 genomic regions, corresponding to 6,605 genes. Remarkably, 2,468 of these genomic regions, corresponding to 15,66 genes, maintained increased accessibility for at least 30 days post-infection (**Figure 4a; Figure S5a, b**). The increased accessible regions (IARs) identified in stNK cells at both 6 and 30 days post-infection were enriched in inflammation regulatory pathways (adjust p<0.05), including the phosphoinositide 3-kinase (PI3K)-Akt, MAPK, Ras, JAK-STAT, and NF-κB signaling pathways (**Figure 4b**). Notably, the NK cell-mediated cytotoxicity pathway was also significantly enriched (**Figure 4b**).

**Figure 4.**
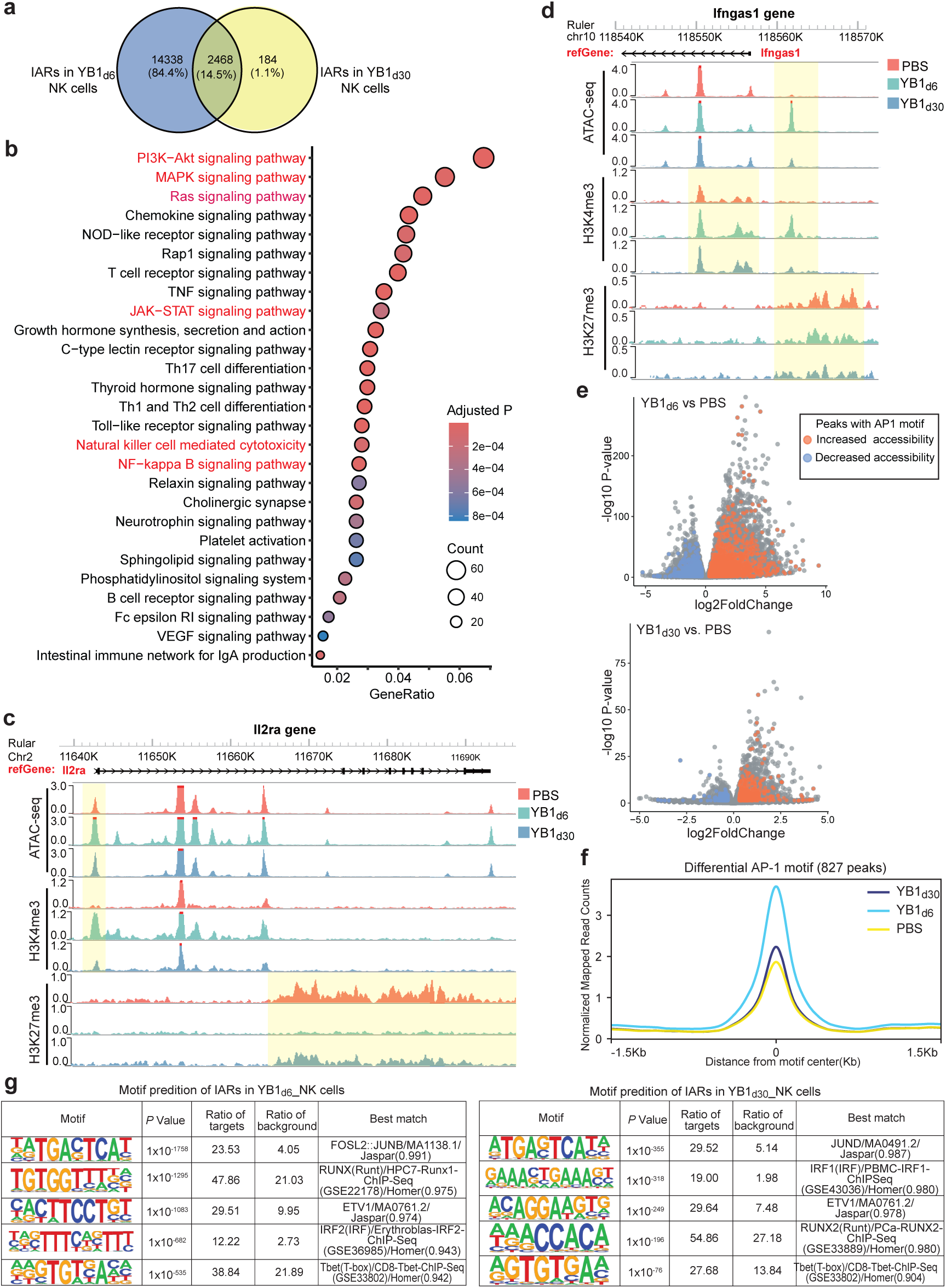
Attenuated *Salmonella* induces long-term epigenetic reprogramming characteristic of trained immunity in NK cells. **a)** Venn diagram illustrating the overlapping increased accessible regions (IARs) in stNK cells on days 6 and 30 post-infection, in comparison to the PBS control group (n = 3 per group, adjusted p-value <= 0.05 and log2 fold change ≥ 0.3). **b)** Dot plot displays enriched signaling pathways and NK cell cytotoxicity-related GO terms for overlapping IARs in stNK cells on days 6 and 30 post-infection. **c-d)** Genome browser tracks of interested gene regions in control NK cells and stNK cells (6 days and 30 days post-infection). **c)** The shadow boxes indicate an increased chromatin accessibility peak and enhanced H3K4me3 marks in the promoter region of the *Il2ra* gene, as well as decreased suppressive H3K27me3 marks in the gene body of *Il2ra*. **d)** The shadow boxes indicate an increased chromatin accessibility peak, enhanced H3K4me3 marks, and a decreased level of suppressive H3K27me3 marks in the promoter region of the *Ifngas1* gene. **e)** Volcano plots display differential chromatin accessibility regions in stNK cells (two time points) compared to control NK cells. Significant peaks with AP-1 motif were colored. **f)** Normalized coverage of overlapped differential AP-1 related accessibility regions between different time points. **g)** Enriched de novo motifs predicted by HOMER among differential accessibility regions in stNK cells compared to control NK cells.

In parallel with the chromatin accessibility analysis, we also performed the CUT&Tag (Cleavage Under Targets and Tagmentation) assay to decipher the histone modification alterations under these conditions, using antibodies against H3K4me3 (active promoters), H3K27me3 (repressive state), and H3K27ac (active enhancers). In the stNK cells, enhanced H3K4me3 marks and lower H3K27me3 marks in the promoter region of the *Kit* gene were detected, together with the multiple prolonged IARs across the whole *Kit* gene body, which indicated higher transcriptional accessibility of the *Kit* gene (**Figure S5c**). Notably, activation of the *Kit*-encoded c-Kit receptor tyrosine kinase has been reported to activate several downstream signal transduction pathways, including PI3K-Akt, MAPK, Ras, and JAK-STAT pathways, which ultimately promote cell survival, proliferation, differentiation, and migration[27]. *Pik3cb*, which encodes an isoform of the catalytic subunit of PI3K involved in the activation of the PI3K-Akt pathway, also showed a coordinated chromatin environment at the promoter region that favored its active expression in the stNK cells (**Figure S5d**). Furthermore, the loci of genes such as *Jak2*, *Stat3*, and *Stat1*, also presented prolonged IARs and histone active patterns in the stNK cells, supporting the activation of JAK-STAT pathway (**Figure S5e-g**).

One of the main memory features of MCMV-specific NK cells and CIML NK cells is the sustained up-regulation of *Il2ra* (CD25, the high affinity IL-2 receptor α chain) in response to IL-12 and IL-18 stimulation, allowing NK cells to be sustained under low physiological levels of IL-2[28, 29]. Similarly, stNK cells showed prolonged epigenetic features that promote *Il2ra* transcription, including increased chromatin accessibility and enhanced H3K4me3 marks in the promoter region, as well as a lower level of suppressive H3K27me3 marks across the gene body (**Figure 4c**). In addition, we found that *ifngas1*, which encodes a long non-coding RNA that promotes IFN-γ expression[30], maintains an active chromatin state that favors its transcriptional accessibility in the stNK cells, as evidenced by increased chromatin accessibility, higher H3K4me3 and lower H3K27me3 marks at its promoter region (**Figure 4d**). Other immune effector genes, such as *Csf2* which encodes the GM-CSF and *Gzmk* which encodes the proteolytic granzyme K, also showed reprogrammed epigenetic alterations favoring their active transcription (**Figure S5h, i**).

Many transcription factors (TFs) play critical roles in the maturation and differentiation of NK cells. The enrichment of JAK-STAT and NF-κB signaling pathways based on specific IARs in the stNK cells suggested that TFs such as STATs and NF-κB could regulate the early initiation of the trained program in NK cells following YB1 treatment (**Figure 4b**). In addition, an analysis of database recorded TF binding motifs in IARs from stNK cells post-infection indicated pronounced enrichment of the AP-1 binding (**Figure 4e, f**). Further de novo TF binding motif prediction from the same IARs in the stNK cells consistently identifies canonical AP-1 motifs (FOSL2/JUNB and JUND separately for two time points) as the most prominent hits (**Figure 4g**). These findings showed that long-term opening of AP-1 motif-containing regions is the core feature of the chromatin landscape changes in the stNK cells and indicated that AP-1 may play a pivotal role in the orchestration of such sustained open chromatin configurations. Notably, it was also observed that the inflammatory memory signature in HCMV-specific NK cells is enriched for AP-1 motifs[31], suggesting the important role of AP-1 in orchestrating the epigenome remodeling in NK cells.

Previous studies have demonstrated that during viral infection, particularly cytomegalovirus (CMV) infection, engagement of cell-surface receptors transmits activating signals downstream to AP-1 through the PI3K-Akt, MAPK, and Ras signaling pathways, ultimately leading to the modification of the epigenome landscape and the formation of inflammatory immune memory in NK cells [26]. Considering the stNK cells displayed many epigenetic features similar to those observed in CMV-specific NK cells, we postulate that the regulatory networks responsible for immune memory formation in NK cells during viral infection may also govern the induction of trained immunity in NK cells following *Salmonella* YB1 treatment.

### Transcriptomic characterization of the NK cell remodeling process after the attenuated *Salmonella* treatment

Given that the epigenome remodeling was initiated during the early acute infection phase and persisted in the latter fully resolution phase, profiling of the entire NK cell training process *in vivo* may provide valuable insights for exploiting these mechanisms therapeutically. We first conducted bulk RNA sequencing of lung-infiltrating NK cells during the acute infection phase (**Figure S6a**). Interestingly, PCA clustering analysis showed that the NK cells at 3 hours post-YB1 infection not only displayed the greatest distance from all other groups, but also locate in the opposite direction along the PC1 axis relative to the PBS-treated NK cells when compared with NK cells at other time points post-YB1 treatment (**Figure S6b**). Considering the rapid decline of NK cells by 3 hours and the obvious recovery by 2 days post-YB1 treatment (**Figure 3a**), it is likely that NK cells at 3 hours post-YB1 treatment have experienced drastic transcriptional changes and may be replaced by the later time points NK cells. Moreover, NK cells on day 1 to day 3 post-YB1 treatment gradually diverged from those in the PBS-treated group, and there is a nearly complete overlap of NK cells on day 2 and day 3 post-YB1 treatment, which is consistent with our previous phenotypic experiment result that the gradually replenished NK cells already exhibit enhanced anti-tumor activity 2 days after YB1 treatment (**Figure 3a-c**). Further analysis of all differentially expressed genes (DEGs) from day 1 to day 5 post-infection revealed that two gene clusters were upregulated during this phase. The first gene cluster was mainly related to cell proliferation and division, while the second one was closely associated with immune defenses (**Figure S6c**). *Zbtb32*, which encodes a TF known for controlling the proliferative burst of MCMV-specific NK cells in response to MCMV infection[32], was significantly up-regulated after YB1 treatment, which explains the boosted proliferation following YB1 treatment (**Figure S6d**). Additionally, most effector cytokine genes (*Ifng* and *Tnf*) and cytotoxic molecule genes (*Gzmb, Gzmc, Gzmk, Gzma, Gzmf,* and *Prf1*) for NK cells’ immune defense function were also significantly upregulated after YB1 treatment (**Figure S6e**).

To further resolve the comprehensive dynamics of immune responses and the intricate NK cell training process following YB1 infection from the acute phase to the resolution phase, we performed single-cell RNA sequencing (scRNA-seq) analysis of lung-infiltrating immune cells (**Figure S7a**). A total of 73,752 high-quality cells encompassing 16 cell types were harvested (**Figure S7b, c**). As a crucial component of the immune defense against bacterial infections, neutrophils rapidly increased in proportion, reaching the peak at 3 hours post-YB1 treatment, followed by a gradual decline (**Figure 5a**). Meanwhile, the proportion of monocytes gradually increased and reached its peak on day 6 post-YB1 treatment (**Figure 5a**). Given that NK cells were conferred with long-lasting anti-metastatic immunity upon YB1 treatment, we extracted the NK cells from the dataset and conducted unbiased re-clustering, identifying four distinct subsets (**Figure S7d; Figure 5b**). Specifically, the expression of *Itgam* and *Cd27* was used to define immature and mature NK cells [33]. Proliferating NK cells were identified through high expression of cell cycle and proliferation hallmarks, *Mki67* and *Birc5*[34]. Finally, exhausted NK cells were identified by high expression of TGF-β/SMAD genes *Smad3/Smad7*, and decreased expression of effector genes such as *Ifng* and *Gzmb*[35]. Consistent with the peripheral NK cell population dynamics revealed by flow cytometry (**Figure 3a**), the proportions of total NK cells as well as their four subsets experienced a sudden decline at 3 hours, followed by a gradual recovery that peaked on day 3 (**Figure 5a-c**). The dynamic changes in immune cell populations during the acute infection phase gradually returned to baseline levels by at least 30 days post-infection, confirming that the resolution of the anti-infectious inflammatory was completed by this time (**Figure 5a-c**). The high expression of most effector molecules in the immature/mature/proliferating NK cell subsets were sustained throughout the acute infection phase and subsequently returned to the baseline levels during the resolution phase (**Figure 5d**). Compared to the other three NK cell subsets, exhausted NK cells consistently exhibited lower expression levels of effector molecules throughout the whole process (**Figure 5d**), aligning with reported functionally impaired phenotype [36]. We further characterized the functions of different NK cell subsets by comparing their pathway activities. During the acute infection phase, most immune defense-related pathways were upregulated, such as the interferon alpha/gamma response, inflammatory response, NK cell activation, TNF-α signaling through NF-κB pathway, MYC targets pathways and IL-2-STAT5 signaling pathway. These changes were predominantly observed in immature NK cells and proliferating NK cells, while mature NK cells showed relatively modest alterations (**Figure 5e**). Notably, even at 30 and 60 days after YB1 treatment, gene ontology terms, including NK cell-mediated tumor cytotoxicity and NK cell cytokine production, remained in slightly more activated states compared to the PBS group (**Figure 5e**). In support of this, a moderate increase in the expression of effector genes *Prf1* and *Gzma* in mature stNK cells 60 days post-infection was detected in the DEGs (**Figure 5f**). The upregulation of TFs, such as AP-1 (*Fosl1* and *Jund*), STATs (*stat1* and *stat5a*) and NF-κB, occurred during the early stages of infection (**Figure 5g**), which supports the predictive conclusion from ATAC-seq analysis that these TFs may orchestrate the establishment of the YB1-induced anti-metastatic epigenetic landscape (**Figure 4b, e-g**). Notably, the expression levels of most early upregulated TFs upon YB1 infection gradually declined, however, certain factors such as *Irf8* and *Stat1* which had been reported to be crucial for the memory formation of MCMV-specific NK cells[37, 38], remained persistently elevated compared to the control samples (**Figure 5g**), further supporting the established NK trained immunity.

**Figure 5.**
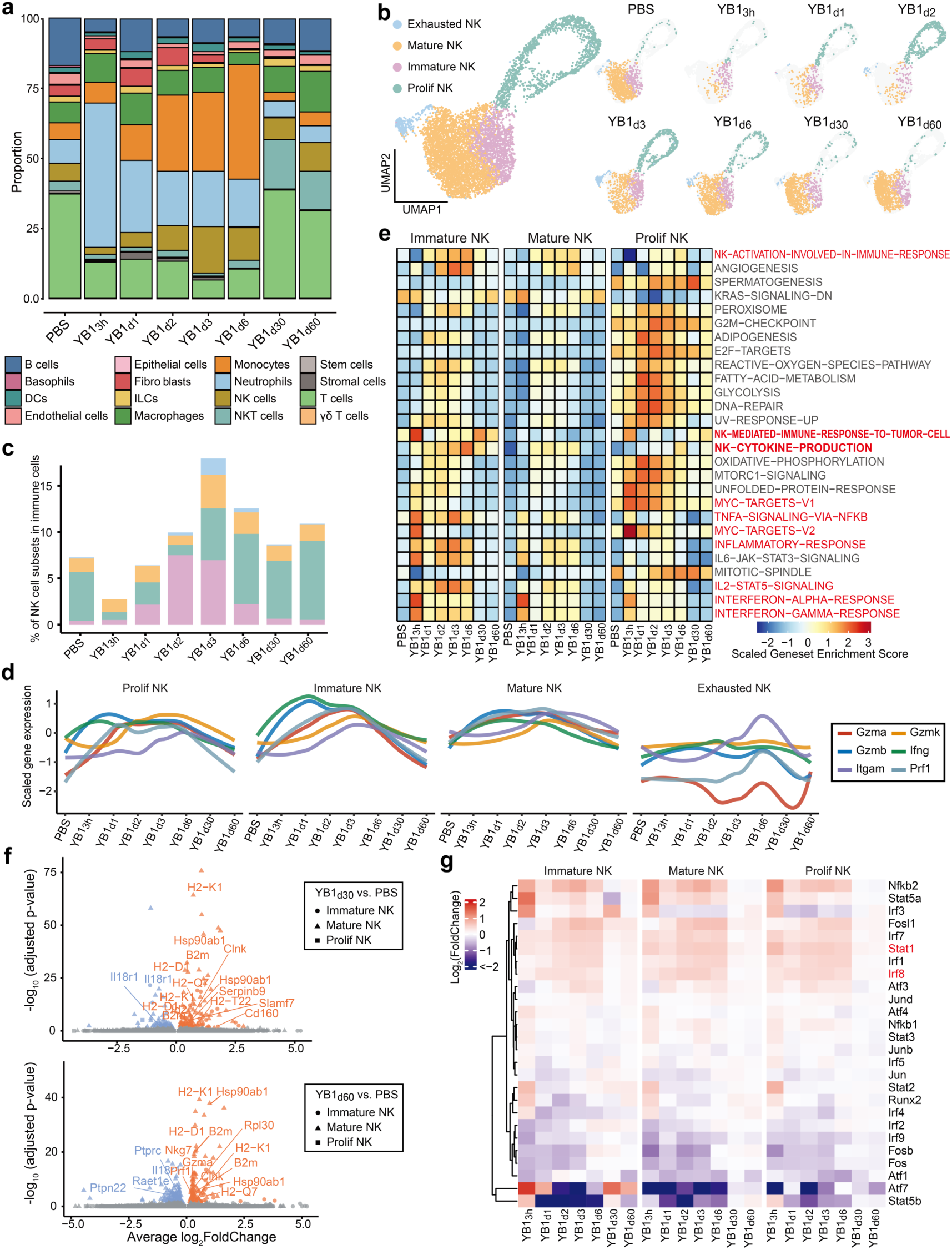
Transcriptomic characterization of the NK cell remodeling process after the attenuated *Salmonella* treatment. A schematic representation for scRNA-seq analysis of lung-infiltrating immune cells is shown in **Figure S7a**. **a)** Stacked bar charts display the dynamic changes of different cell populations across different groups. PBS treatment serves as a control. **b)** UMAP shows subsets of NK cells. **c)** Stacked bar plot shows the percentage of different NK cell subsets to total immune cells across different groups. **d)** Normalized gene expression patterns of NK cell maturation marker *Itgam* and cytotoxic effector genes along the timeline post-YB1 treatment in different NK cell subsets. **e)** Activities of different hallmark pathways and NK cell-related gene ontology terms across different groups. **f)** Volcano plots showing differentially expressed genes in stNK cells on day 30 or 60 post-YB1 treatment compared to the PBS group. Significantly differentially expressed genes are colored (adjusted p-value ≤ 0.05 and absolute log2 fold change ≥ 0.3), and different NK cell subtests are indicated as different shapes. **g)** The heatmap displays the activity of different transcription factors, with color coding representing the log2 fold changes of the average AUC score in different YB1-treated groups compared to the PBS group. Only AP-1, STATs, and NFkB-related regulons were shown.

Taken together, our bulk and single-cell RNA-seq analyses of NK cells revealed dynamic population shifts and transcriptomic reprogramming. These data strongly indicate that inflammatory signaling pathways activated in the potentially trained NK cells — particularly immature and proliferating subsets — play pivotal roles in establishing trained immunity.

### IL-12 and IL-18 are indispensable for the development of stNK cells

Cytokine regulation plays a crucial role in the development, differentiation, and effector function of NK cells. Pro-inflammatory cytokines, such as IL-2, IL-12, IL-15, IL-18, IL-33, IFN-γ, and type I IFNs, have been reported to be instrumental in forming antigen-specific or non-antigen-specific memory in NK cells. Among these cytokines, the combination of IL-12 and IL-18, along with homeostatic cytokine IL-15, is particularly effective in promoting changes in chromatin accessibility[26]. Notably, these three cytokines are mainly produced by other immune cells. Considering the significant responses of stNK cells to IL-12 and IL-18 revealed by our ATAC-seq analysis (**Figure 4c**), we conducted investigation into the interactions between other immune cells and NK cells, especially interactions that are mediated by IL-12/IL-15/IL-18 and their receptors. During the acute infection phase, neutrophils, monocytes, and macrophages constituted the primary NK cell-interacting populations (**Figure S8a)**, exhibiting significantly enhanced interactions with NK cells orchestrated by IL-12/IL-15/IL-18 and their receptors (**Figure 6a**). Additionally, these three myeloid cell types are the most abundant immune cell types following YB1 treatment (**Figure 6b**). We thereafter quantified the levels of 111 cytokines and chemokines (including IL-12 and IL-15) in murine serum at various time points following YB1 treatment (**Figure 6c**; **Figure S8b, c**). The quantification showed a rapid surge of systemic IL-12 production at 3 hours post-YB1 treatment, which became undetectable 1 day later (**Figure 6c**). The quantification also revealed a slight increase of IL-15 on day 1 and day 2 post-YB1 treatment (**Figure S8c**). Regarding IL-18, our previous study showed that systemic IL-18 levels peaked rapidly by 3 hours post-YB1 infection, gradually returning to baseline within one week[18]. Of note, IL-2, a crucial cytokine for promoting NK cell proliferation and cytotoxicity[39], exhibited low abundance and remained unchanged following YB1 treatment (**Figure 6c; Figure S8c**). Collectively, the cytokine quantification suggests that the NK cell population experienced short-term exposure to a high level of IL-12 (within one day post-infection), minor increased exposure to IL-15, and prolonged exposure to high levels of IL-18 (around 1 week) during the early initiation of the trained program following YB1 treatment.

**Figure 6.**
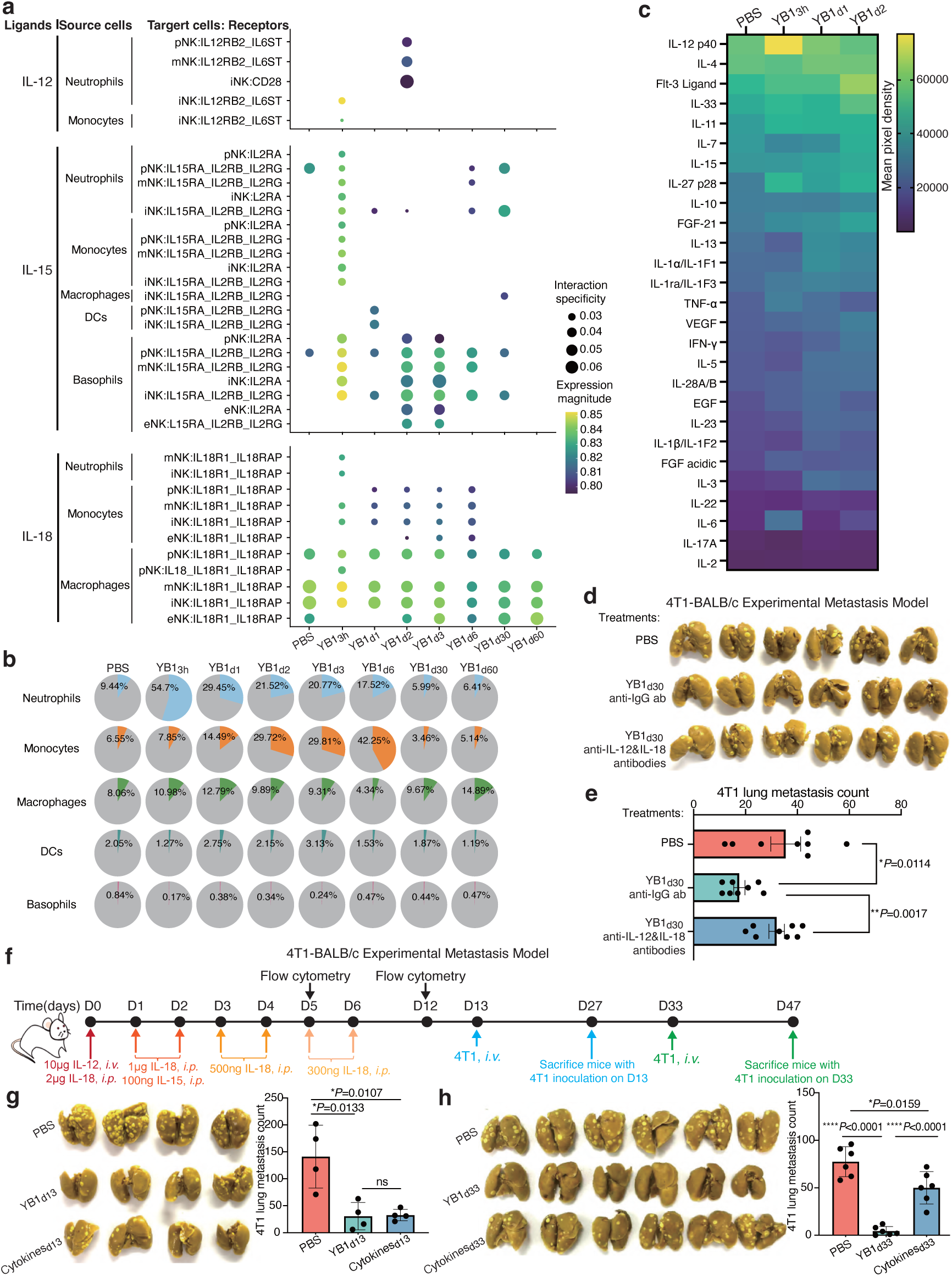
IL-12 and IL-18 are indispensable for the development of stNK cells. **a)** Bubble heatmap displays interaction intensity between myeloid cells and NK cells through stimulation ligands IL-12, IL-15, and IL-18 produced by myeloid cells to relevant receptors on different NK cell subsets (pNK: Prolif NK, iNK: Immature NK, mNK: Mature NK, eNK: Exhausted NK). **b)** Proportion dynamics of different myeloid cell types among total immune cells within the PBS and YB1 treatment groups. **c)** The cytokine array heatmap displays the dynamics of selected cytokines and chemokines in the serum of mice treated with PBS or YB1 at various time points post-infection. **d-e)** The picture of lungs fixed in Bouin solution (**d**) and the quantification (**e**) of 4T1 lung metastases (n=8 per group) from BALB/c mice received different treatments: PBS, YB1, or YB1 plus IL-12 and IL-18 neutralization. **f)** A schematic representation showing the experimental design to simulate the *in vivo* IL-12/IL-15/IL-18 dynamics induced by YB1 treatment. PBS- or YB1-treated mice served as controls. **g)** The picture of lungs fixed in Bouin solution and the quantification of 4T1 lung metastases (n=4 per group, shown one representative result of 2 independent experiments, presented as mean values ± SD) for the BALB/c mice received 4T1 cancer cell inoculation on day 13 as depicted in Figure 6f. **h)** The picture of lungs fixed in Bouin solution and the quantification of 4T1 lung metastases (n=6 per group) for the BALB/c mice received 4T1 cancer cell inoculation on day 33 as depicted in Figure 6f. Combined results of 2 independent experiments are displayed and presented as mean values ± SEM for Figure 6 **e, h**. All *P* values are yielded by two-tailed unpaired t-tests and are shown in the relevant figure.

To confirm the necessity of high levels of IL-12 and IL-18 in the training of NK cells upon YB1 treatment, we conducted *in vivo* neutralization of these cytokines using specific antibodies and evaluated their effects on the YB1-induced long-term anti-metastatic capability (**Figure S8d**). The results showed that *in vivo* neutralization of IL-12 and IL-18 significantly abolished the long-term anti-metastatic effects of YB1 treatment (**Figure 6d, e**). Subsequently, we attempt to simulate the *in vivo* dynamics of IL-12/IL-15/IL-18 induced by YB1 treatment (**Figure 6f**). To achieve this, each mouse received only one high dose of IL-12 (via *i.v.* injection) on day 0, followed by low doses of IL-15 (via *i.p.* injection) on days 1 and 2. Additionally, starting from day 0 until day 6, each mouse received a high dose of IL-18 with gradually decreasing amounts every day (via *i.p.* injection). Mild ruffled fur, which is a common symptom observed in infected mice such as YB1-treated mice, was also observed in the mice injected with cytokines (**Figure S8e**). Cytokine injections stimulated the expansion of NK cells *in vivo* within 5 days, which returned to the baseline levels as in uninfected PBS group on day 12, resulting in a similar population dynamic as that induced by YB1 treatment (**Figure S8f**). To separately validate the effects of cytokine injections on short-term and long-term anti-metastatic ability, we injected 4T1 cancer cells into cytokine-treated mice on either day 13 or day 33, with PBS- or YB1-treated mice serving as controls (**Figure 6f**). Cytokine injections resulted in a comparable short-term anti-metastatic ability to YB1 treatment (**Figure 6g**). However, the cytokine-treated mice failed to exhibit long-lasting anti-metastatic ability as in the YB1-treated mice (**Figure 6h**). These findings collectively indicate that exposure to IL-12/15/18 alone enables the expansion and activation of NK cells for a short period but is insufficient to mount long-term anti-metastatic immunity in NK cells.

### Anti-metastatic trained immunity induced by attenuated *Salmonella* outperforms immune checkpoint blockade therapies in suppressing metastasis

Immune checkpoint blockade therapy, especially PD-1 blockade therapy, represents one of the most significant breakthroughs in the field of cancer immunotherapy and has been reported to prevent metastasis in mice and humans [19, 40]. To compare the anti-metastatic effects of the stNK cells with immune checkpoint blockade, we applied these two immunotherapies to an experimental metastasis model established with 4T1/4T1-Luci cancer cells and BALB/c mice. Compared to the potent anti-metastatic ability induced by YB1 treatment, PD-1 blockade only displayed a limited effect in reducing 4T1 lung metastatic nodules (**Figure S9a; Figure 7a, b**), and showed no effect in inhibiting the early survival of 4T1-Luci cancer cells in the lung by *in vivo* imaging analysis (**Figure S9b; Figure 7c, d**). Apart from PD-1, TIGIT is another well-known immune checkpoint molecule modulating NK cell and T cell activation [20, 41]. Compared to the potent anti-metastatic effect induced by YB1 treatment 6 days or 30 days post-infection, TIGIT blockade showed no effect in suppressing metastasis (**Figure 7e, f; Figure S9c-e**). These data collectively suggest that the anti-metastatic trained immunity induced by attenuated *Salmonella* YB1 outperforms immune checkpoint blockade therapies in suppressing metastasis, highlighting the importance and potential of inducing trained immunity in NK cells as an effective anti-metastatic strategy with long-term effects.

**Figure 7.**
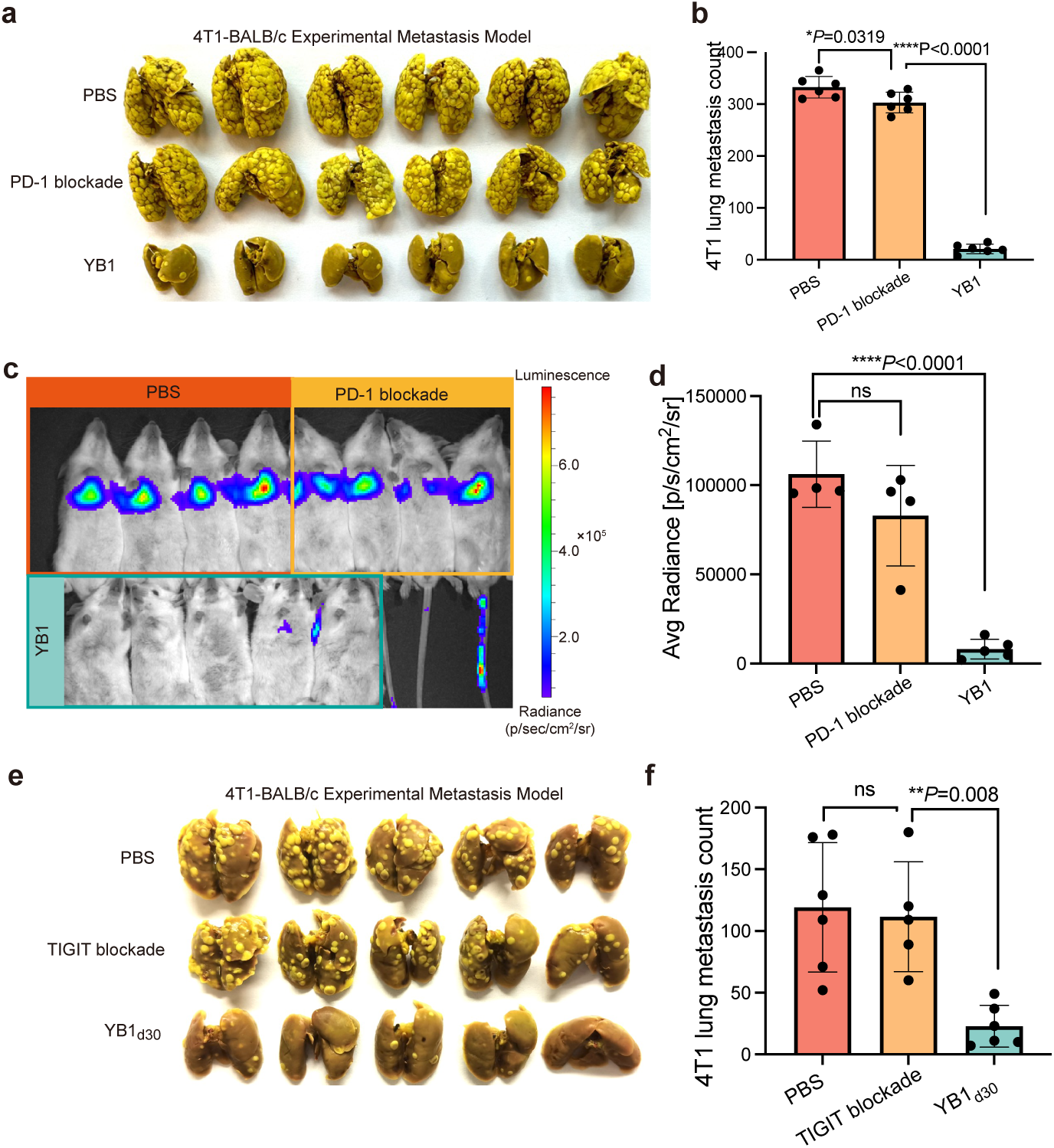
Anti-metastatic trained immunity induced by attenuated *Salmonella* outperforms immune checkpoint blockade therapies in suppressing metastasis. **a-b)** Images of Bouin-fixed lungs and the quantification of 4T1 lung metastases (n=6 per group) for the mice received different treatments: PBS, PD-1 blockade, and YB1, as depicted in **Figure S9a**. **c-d)** Tracking and quantification (n = 4 for PBS and PD-1 blockade groups, n=5 for YB1) of 4T1-Luci cells *in vivo* using luciferase live imaging in differently treated BALB/c mice, as depicted in **Figure S9b**. **e-f)** Images of Bouin-fixed lungs (**e**) and the quantification (**f**) of 4T1 lung metastases (n=6 for PBS and YB1_d30_, n=4 for TIGIT blockade group) for the mice received different treatments: PBS, TIGIT blockade, and YB1_d30_. The experimental schedule is shown in **Figure S9d.** One representative experiment of two independent experiments is displayed and presented as mean values ± SD for **Figure 7b, d, and f**. All *P* values are yielded by two-tailed unpaired t-tests and are shown in the relevant figure.

## Discussion

Trained immunity, an evolutionarily ancient program of immunological memory that results in enhanced reactions to secondary stimulation, has been extensively demonstrated in myeloid cells as a host defense mechanism against heterologous pathogen infections. This process can be triggered by various stimuli, such as the BCG vaccine, *Candida albicans*, influenza, β-glucan, CpG, and LPS [2–6]. Notably, the phenomenon of immune tolerance, characterized by a reduced immune function after restimulation, can occur as the opposite of trained immunity, particularly following treatment of myeloid cells with LPS or specific microbial pathogens, such as *Candida albicans* (high dosage), influenza A virus, and *Escherichia coli* (*E. coli*)[2, 42–44]. The nature and dose of the immune stimulus are critical in determining whether trained immunity or tolerance is induced[45]. Despite these intriguing findings, the functional effects of inducing trained immunity with appropriate stimuli in innate lymphoid cells other than in myeloid cells have not been thoroughly investigated, especially in the scenario of prevention of cancer progression and metastasis. In this study, we showed that *Salmonella* YB1 is an unreported stimulus that can induce long-lasting anti-metastatic trained immunity in NK cells, an underappreciated cell type in current trained immunity research. Our study broadens the scope of the trained immunity field by expanding the range of stimuli, the types of immune cells being trained, and the functional scenarios. This research also underscores the potential of harnessing trained immunity in NK cells as a long-lasting and effective antitumor strategy.

Two studies have suggested that inducing trained immunity in macrophages can be beneficial for long-term anti-metastatic responses. In one study, intranasal infection with the influenza virus has been found to train respiratory mucosal-resident alveolar macrophages to enhance their tumor cell phagocytotic ability, resulting in long-lasting and tissue-specific anti-metastatic immunity[9]. In the other study, the induction of pneumonia-related sepsis through intratracheal administration of *E. coli* leads to the training of alveolar macrophages, which then exert an anti-metastatic effect by promoting the anti-metastatic activity of tissue-resident T cells, particularly the γδ T cells[10]. Unlike the trained alveolar macrophages in these studies, our study showed that trained NK cells, upon treatment with *Salmonella* YB1, directly exert a long-term anti-metastatic effect by eliminating cancer cells. Of note, during the acute infection phase, various innate immune cells, such as neutrophils and monocytes, are sequentially upregulated as host defense mechanisms against *Salmonella* YB1 (**Figure 5a**), and these myeloid cells secrete essential cytokines involved in the early training of NK cells (**Figure 6a, b)**. Therefore, it cannot be excluded that myeloid cells contribute to the anti-metastatic effects induced by *Salmonella* YB1 via promoting the early initiation of the trained program in NK cells. Moreover, the anti-metastatic effects of *Salmonella* YB1 are systemic, whereas the effects of influenza virus and *E. coli* in the two previous studies are lung-specific, which are probably attributed to their different routes of administration. Collectively, our study unveils a novel long-term anti-metastatic trained immunity exerted by NK cells rather than by myeloid cells or adaptive immune cells.

Different from the tumor cell recognition by cytotoxic T cells, which requires MHC-I-restricted antigen presentation by cancer cells, NK cells do not rely on neoantigens or tumor-associated antigens to recognize tumor cells. More importantly, tumor cells with low MHC-I expression levels are naturally more vulnerable to NK cell killing[46]. While NK cells’ activity was initially considered “spontaneous”, it is now appreciated that NK cells require an array of signals to be adequately primed for optimal effector potential. Analogous to developing immune memory in T cells, the formation of antigen-specific immune memory in NK cells requires integrating antigen-specific signaling, co-stimulation signaling, and cytokine signaling[26]. MCMV-specific NK cells, for instance, develop immune memory, manifested as enhanced IFN-γ production and cytotoxicity, by integrating signals from antigen-specific receptor Ly49H[47], costimulatory molecule DNAM1[48], and cytokines (IL-12/IL-18/IL-33)[49–51]. Although *in vitro* cytokine-primed CIML NK cells exhibit both higher IFN-γ production and stronger cytotoxicity against tumor cells while they are still in a stimulated active state[52], they only maintain the potential for enhanced IFN-γ production but not for enhanced cytotoxicity upon returning to homeostasis [14, 15], suggesting that additional signals are required to mount the optimal immune memory consisting of both enhanced cytokine production and cytotoxicity potential. This hypothesis is supported by another study demonstrating that adoptive transfer of CIML NK cells into irradiated mice immediately after irradiation (3 hours) - but not into non-irradiated mice - resulted in suppression of established RAM-s tumor growth in a manner dependent on both IFN-γ production and perforin-mediated cytotoxicity, without either of which the inhibitory effect could not be achieved[29]. These findings also explain why simulating the *in vivo* dynamics of IL-12/IL-15/IL-18 induced by *Salmonella* YB1 treatment provides only transient anti-metastatic effects, unlike the sustained anti-metastatic response generated by the treatment itself (**Figure 6f-h**). Therefore, additional signals beyond pro-inflammatory cytokines like IL-12 and IL-18 are likely required to induce long-lasting anti-metastatic immune memory in NK cells following *Salmonella* YB1 treatment. Identifying these signals will be crucial for further understanding and harnessing NK cell-mediated trained immunity.

It is widely recognized that 90% of cancer-related deaths result from cancer metastasis, and novel preventive and therapeutic strategies/agents specifically targeting metastasis remain in urgent need[53]. Considering that stNK cells induced one month before (**Figure 1l-n**) or one week after[18] orthotopic inoculation of 4T1 breast cancer cells significantly reduced post-surgical lung metastasis in the breast cancer spontaneous metastasis model, our study provides insight into the prophylactic induction of anti-metastatic trained immunity in NK cells prior to surgical resection of newly diagnosed primary tumors in clinical settings. This could be achieved either by applying the attenuated *Salmonella* YB1 strain as a bacterial cancer vaccine or by adoptive transfer of stNK cells generated *in vitro* in the future. Such a strategy is particularly promising for cancer types with high risks of metastasis.

## Experimental Section

### Animals

The BALB/c, C57BL/6J, Nude (BALB/cAnN-nu), and NSG (NOD.Cg-Prkdc^scid^Il2rg^tm1Wjl^/SzJ) mice, aged 6 to 8 weeks, were obtained from the Laboratory Animal Unit of Shenzhen Institutes of Advanced Technology or The University of Hong Kong. The B6 *Rag1* knockout mice (JAX stock # 034159, C57BL/6J-Rag1^em10Lutzy^/J mice) and B6 TCR-delta knockout mice (B6.129P2-*Tcrd*^tm1Mom^/J) were purchased from the Jackson Laboratory and housed in the Laboratory Animal Unit of The University of Hong Kong. The CD45.1 transgenic mice (Pep Boy mice) were obtained from Prof. Liwei LU. All mice were housed in a 12 h light/12 h dark cycle at ∼23 °C and 40% relative humidity. All mice experimental protocols were approved by the Committee on the Use of Live Animals in Teaching and Research (CULATR) of The University of Hong Kong, with reference number 5410-20.

### Bacterial strains and cell lines

*Salmonella typhimurium* strain YB1 was genetically engineered in our laboratory[16]. *Salmonella* YB1 was cultured in LB medium supplemented with DAP (100 µg/mL, Sigma D1377-5G), chloramphenicol (25 µg/mL), and streptomycin (50 µg/mL). Mouse 4T1 breast cancer cells and CT26 colon cancer cells were purchased from ATCC. MB49 cancer cells were generously provided by Dr. LIU Chenli from the Shenzhen Institute of Synthetic Biology. Murine YAC-1 cells were kindly provided by Stem Cell Bank, Chinese Academy of Sciences. The 4T1-Luci cell lines were generated using a PiggyBac transposon system carrying firefly luciferase. The 4T1 breast cancer cells, CT26 colon cancer cells, and YAC-1 cancer cells were maintained in RPMI 1640 medium supplemented with 10% fetal bovine serum, streptomycin, and penicillin in a tissue incubator at 37°C and 5% CO_2_. The MB49 bladder cancer cells were maintained in DMEM medium supplemented with 10% fetal bovine serum, streptomycin, and penicillin in a tissue incubator at 37°C and 5% CO_2_.

### Breast cancer orthotopic metastasis model

To establish the 4T1-BALB/c orthotopic metastasis model, 4T1 cells were cultured to 50%-80% confluency, harvested, and washed three times with sterile PBS. The cells were then stained with 0.4% trypan blue to assess cell viability using a hemacytometer. The viable 4T1 cells were diluted to a concentration of 1×10^6^/mL in PBS, and 100 µL of the resuspended cells were injected into the mouse fat pad using a 27G needle. This resulted in the development of a primary tumor in the breast, subsequently leading to spontaneous lung metastasis within one month. Primary tumor size was measured using calipers and was calculated as 4/3 × π × (height × width^2^)/8. Lung metastatic nodules were counted under a stereomicroscope.

### Experimental metastasis models

To establish the breast cancer experimental metastasis model, 1.2×10^5^ 4T1 cells (unless indicated in the schematic representation), resuspended in 100 µL PBS, were intravenously injected into each BALB/c, Nude, or NSG mouse via the lateral tail vein. For bladder cancer experimental metastasis models, 5×10^5^ MB49 cells resuspended in 100 µL PBS were intravenously injected into each C57BL/6J, *Rag1*, or knockout mouse via the lateral tail vein. To establish the liver experimental metastasis model, each BALB/c mouse was injected via portal vein with 1×10^5^ CT26 cells. Metastasis in the above models was quantified by counting lung metastatic nodules under a stereomicroscope or by H&E staining.

### Bacterial treatment

The *Salmonella* used in the treatments were prepared from overnight cultures, and their cell numbers were determined by measuring the optical density at 600 nm (1 OD = 1×10^9^ CFU). The desired concentration of bacteria was achieved by washing them three times in sterile PBS. For experiments using immunocompetent BALB/c or C57BL/6J mice, 2×10^7^ CFU of *Salmonella* YB1 (unless indicated in the schematic representation) were intravenously injected into each mouse through the tail vein. For immunocompromised Nude mice, NSG mice, and *Rag1* knockout mice, 1×10^7^ CFU of *Salmonella* YB1 were injected for each mouse via the tail vein. Administration of PBS serves as the control treatment in comparison to the bacterial treatment.

### Luciferase live imaging

Female BALB/c mice, 8 weeks old, were pretreated with *Salmonella* YB1 on day 0. At different time points following *Salmonella* YB1 treatment, 6×10^5^ 4T1-luci cells were intravenously injected through the tail vein of the BALB/c mice. For *in vivo* imaging, mice were anesthetized with ketamine and medetomidine, and then 100 µL of D-luciferin (30 mg/mL, Gold Biotechnology) was intraperitoneally injected 5 minutes prior to live imaging. Images were captured using the PE IVIS Spectrum *in vivo* imaging system and analyzed using Living Image 4.0 software. The surface intensity of bioluminescence was measured using the region of interest (ROI) tools provided by Living Image 4.0 software.

### Histopathology analysis

Lung or liver tissues were fixed in 4% paraformaldehyde in PBS for 24 hours at 4°C with shaking. Subsequently, the tissues were dehydrated and embedded in a paraffin block, and 5µm-thick sections were prepared for hematoxylin and eosin (H&E) staining.

### NK Cell depletion *in vivo*

To assess the impact of NK cells on the long-term anti-metastatic effect induced by *Salmonella* YB1 in BALB/c mice, *in vivo* depletion of NK cells was initiated on day 29 through intraperitoneal (*i.p.*) injection of 35µL of anti-asialo GM-1 antibody (Cat. no#146002; clone Poly21460; BioLegend). This was followed by additional *i.p.* injections of anti-asialo GM-1 antibody on days 30 (30µL), 33 (30µL), 36 (30µL), 39 (25µL), and 42 (20µL). Matched isotype rabbit polyclonal IgG (Cat. no #BE0095; polyclone; BioXcell) served as the control. In C57BL/6J mice, *in vivo* NK cell depletion was initiated on day29 through intraperitoneal (*i.p.*) injection of 600µg of anti-NK1.1 antibody (Cat. No #BE0036; clone PK136; Bioxcell). This was followed by additional *i.p.* injections of anti-NK1.1 antibody on days 30 (300µg), 31 (300µg), 33(500µg), 36 (300µg), 39 (200µg) and 42 (200µg). Matched isotype mouse polyclonal IgG2a (Cat. no #BE0085; clone C1.18.4; BioXcell) served as the control. Confirmation of NK cell depletion in BALB/c mice was achieved by assessing the absence of CD3– NKp46+ cells in peripheral blood. Similarly, in C57BL/6J mice, the depletion of NK cells was confirmed by evaluating the absence of CD3– NK1.1+ cells in peripheral blood.

### Flow cytometry

For the detection of cell surface markers, single-cell suspensions were incubated with antibodies for 30 minutes on ice, followed by two washes with 1% BSA/PBS before flow cytometric analysis. In selected experiments, propidium iodide (PI) with a final working concentration of 1 µg/mL or DAPI with a final working concentration of 500 ng/mL was added to the cell suspension immediately prior to flow cytometric analysis to gate viable cells. For the intracellular detection of IFN-γ, immune cells were stimulated *ex vivo* with or without PMA/ionomycin in the presence of brefeldin A for 5 hours, followed by cell surface staining as mentioned above. Subsequently, cells were fixed and permeabilized using the BD Fixation/Permeabilization kit (catalog number 554714), following the manufacturer’s instructions. The cells were then stained with an anti-IFN-γ antibody on ice for 30 minutes and washed twice prior to flow cytometric analysis. Flow cytometric events were acquired using either the ACEA Novocyte Flow Cytometer or the BD FACSAria™ Fusion Flow Cytometer. FlowJo_v10.6.2 and NovoExpress_v1.4.1 software were used for the analysis of the flow cytometric data. Antibodies for flow analysis were summarized in Table S1.

### Isolation of lung-infiltrating immune cell

To obtain single-cell suspensions from the lung tissue, minced tissue chunks were incubated with Collagenase Type IV (Sigma-Aldrich, catalog number C5138) and DNase I Type IV (Sigma-Aldrich, catalog number D5025) in RPMI1640 medium at 37°C for 60 minutes. Upon completion of this incubation, red blood cells were lysed, and the resulting single-cell suspension was resuspended in 3 mL of 40% Percoll and carefully layered onto 3 mL of 70% Percoll. Immune cells were subsequently enriched from the interface between the 40% and 70% Percoll layers following density gradient centrifugation at 1260 × g for 30 minutes at room temperature.

### Degranulation assay

Lung-infiltrating immune cells were co-cultured with or without 1× the number of YAC-1 cells for 5 h at 37℃ in 5% CO2. GolgiStop (BD) and anti-CD107a (BD Biosciences, catalog no. 560648) were supplemented during the culture. After culture, cell mixtures were harvested and stained for cell surface markers and subjected to flow cytometric analysis.

### Purification and culture of murine NK cells

Murine lung-infiltrating immune cells were isolated from the lung by density gradient centrifugation with 40% and 70% Percoll. Murine NK cells were purified using the NK Cell Isolation Kit (Miltenyi Biotec, catalog no.130-115-818) according to the manufacturer’s instructions. Purified NK cells were resuspended in Iscove’s modified Dulbecco’s medium (IMDM) supplemented with 10% fetal calf serum, L-glutamine (1 mM; Gibco), streptomycin (100 µg/mL; Sigma), penicillin (100 IU/mL; Sigma) and 50 µM beta-mercaptoethanol.

### Cytotoxicity assay

Effector NK cells were co-cultured with target cells, which have been labeled with CellTrace Violet (ThermoFisher, catalog no.C34571), in a 96-well plate at various effector: target (E/T) ratios for 5 hours at 37°C. Following the co-culture period, cell mixtures would be harvested and washed prior to flow cytometric analysis. To determine cell viability, target cells that exhibit positive staining for PI will be considered as deceased cells, allowing for the calculation of the percentage of dead cells. The percentage of specific lysis was calculated for each well as follows: % of specific lysis =100 × (Dead Ratio^experimental^ -Dead Ratio^spontaneous^)/ (Dead Ratio^maximum killing^ - Dead Ratio^spontaneous^).

### Bone marrow transplantation

Bone marrow cells were extracted from C57BL/6N (CD45.2+) mice 23 days after PBS or YB1 treatment and transplanted into naïve, sub-lethal irradiated (850cGy), syngeneic Pep boy (CD45.1+) mice. Successful reconstitution can be achieved at 5 weeks post- bone marrow cell transfer.

### Bulk RNA-sequencing and data analysis

For each replicate, approximately 30,0000 viable NK cells were sorted and subsequently lysed in TRIzol reagent (Thermo Fisher), followed by RNA extraction. PolyA-positive RNA was isolated by Dynabeads™ mRNA Purification Kit (Thermofisher) and subjected to mRNA library preparation according to NEBNext Ultra RNA Library Prep Kit for Illumina (New England Biolabs) following manufacturer’s recommendations. Finally, paired-end sequencing was performed on an Illumina NovaSeqPE150 platform (Novogene). Gene expression profiles of each sample were evaluated by Salmon[54]. Significant differential expressed genes (DEGs) were acquired with the help of DESeq2[55]. The hierarchical clustering of DEGs was based on their normalized expression, and the number of gene clusters was specified manually. Gene ontology enrichment analysis was carried out by ClusterProfile[56].

### ATAC-seq library preparation

The ATAC-seq library was prepared using the refined Omni-ATAC procedure to minimize contamination from mitochondrial DNA[57]. In brief, 50,000 viable NK cells were sorted and subsequently washed once with cold PBS. The cells were then resuspended in 50μl of cold lysis buffer (10 mM Tris-HCl (pH 7.4), 10 mM NaCl, 3 mM MgCl2, 0.1% IGEPAL CA-630, 0.1% Tween-20, and 0.01% Digitonin). After a 5-minute incubation, the cell lysates were diluted with 1 mL of cold buffer (10 mM Tris-HCl (pH 7.4), 10 mM NaCl, 3 mM MgCl2, 0.1% Tween-20) and subsequently centrifuged at 500g for 10 minutes at 4 °C. The resulting nuclei were resuspended in 50μl of transposition reaction mix (25μl of TD buffer, 2.5μl of Tn5 transposase, and 22.5μl of nuclease-free water; Illumina)), and incubated for 30 minutes at 37 °C. The transposed DNA fragments were purified using a Qiagen MinElute Reaction cleanup kit. Subsequently, the purified DNA was barcoded with Nextera indexes and subjected to PCR amplification for 13 cycles using NEBNext high fidelity 2× PCR master mix (New England Biolabs). PCR products were purified and double-selected using AMPure XP beads before quality control by the Agilent 2100 Bioanalyzer system (Agilent) and sequenced using the Illumina NovaSeqPE150 platform (Novogene).

### CUT&Tag library preparation

The CUT&Tag follows previously reported protocol[58]. Freshly sorted 50,000 NK cells were centrifuged for 5 min at 500g, resuspended in 100 μl antibody buffer (20 mM HEPES pH 7.5, 150 mM NaCl, 4 mM EDTA, 0.5 mM spermidine, 0.05% digitonin, 0.01 % NP-40, 1× protease inhibitors, and 1% BSA) and incubated for 3 min on ice to extract nuclei. Nuclei were then centrifuged at 600g for 3 min, resuspended in 10 μl antibody buffer, and incubated with 10µL of activated Con-A beads. After washing twice with antibody buffer, the nuclei-conA beads complex was resuspended with 50 μl antibody buffer, incubated with 1 μg primary antibodies (Rabbit-H3K4me3 (Abcam ab8580), Rabbit-H3K27me3 (Cell Signaling Technology, cat. no. 9733), Rabbit-H3K27ac (Active Motif Catalog No: 39085)) and control IgG (Rabbit-IgG (Millipore Catalog No: 12-370)) at 4℃overnight. After washing with antibody buffer, 1:500 secondary antibodies were added to bind to the primary antibodies. Then, 1.5 μl 4 μM pA-Tn5 was added to each reaction and incubated at room temperature for 1 hour. The cell nuclei were washed with Dig300 buffer (20 mM HEPES pH 7.5, 300 mM NaCl, 0.5 mM spermidine, 0.01% digitonin, 1× protease inhibitors, and 1% BSA) and resuspended in 100 µL of Tagmentation buffer (Dig300 buffer containing 10 mM MgCl2) and incubated at 37 °C for 1 hour. The genome DNA was then purified with the Qiagen MinElute Reaction cleanup kit. Subsequently, the purified DNA was barcoded with Nextera indexed primers and subjected to PCR amplification using NEBNext high fidelity 2× PCR master mix. The resulting PCR products were purified and double-selected with AMPure XP beads before quality control and sequencing on the Illumina NovaSeqPE150 platform (Novogene).

### Data analysis for ATAC-seq and CUT&Tag-seq

Adapter sequences and low-quality bases were trimmed by fastp[59]. The resulting reads were mapped to the mouse mm10 reference genome with the help of bowtie2[60] in a very sensitive mode. Duplicated reads were marked by sambamba[61] with default parameters. Further filtration was conducted by deepTools[62] including removing reads with mapping quality lower than 30 or reads located in the mouse ENCODE blacklist region[63]. In addition to the above filtration, read shift was applied to ATAC-seq samples. Peaks were called using the MACS3 software suite. Consensus peaks were merged from each sample in the ATAC-seq and CUT&Tag-seq experiments respectively. Reads in consensus peaks were counted using DiffBind[64]. DESeq2[55] was used for differential histone modification or chromatin accessibility between different conditions, signals with p-value less or equal to 0.05 and log2 fold change larger or equal to 0.3 were considered significant. Gene ontology enrichment analysis of significantly increased chromatin accessibility peaks was conducted in clusterProfiler[56] using the nearest gene annotated. JASPAR2020[65] mouse core database was used for motif matching in peaks with the help of motifmatchr[66]. Homer v4.11 findMotifsGenome.pl was used for de novo motif prediction on significantly different chromatin regions [67]. To simplify the plots of epigenome signal visualization, genome browser tracks were created with the bamCoverage command in deepTools[62] on merged bam files from repeats in the same condition, with the binSize set to 5 and normalize using CPM (Counts Per Million mapped reads). Track visualization was implemented by the WashU Epigenome Browser[68].

### Single-cell RNA sequencing (RNA-Seq) and data analysis

At various time points after *Salmonella* YB1 injection, lung-infiltrating immune cells were isolated using a Percoll-based density centrifugation method. Samples were collected from three independent mice at each time point, except for the PBS group, which included data from 6 independent mice. This method resulted in a purity of over 90% for the isolated immune cells. scRNA-seq data was generated with a 10X Genomics Chromium workflow using the Chromium Next GEM Single-Cell 3ʹ Reagent Kit (v3.1, 10X Genomics) and sequenced on Illumina NextSeq 500. Cellranger[69] pipeline was first utilized to align reads to the mouse reference genome (mm10, downloaded from https://support.10xgenomics.com/) and single-cell level gene expression evaluation for each sample. Seurat V5[70] package was used for further procession of scRNA-seq data. Low-quality cells with less than 200 genes expressed or have a high level of mitochondrial genes expressed (mitochondrial genes expression / total genes expression > 10%), and doublets with more than 8,000 genes expressed were filtered out for further analysis. All samples’ data sets were integrated into one Seurat data set based on the anchors identified from the top 30 principal components in each sample. Dimensionality reduction using UMAP and K-nearest neighbor clustering with resolution set to 1 were then applied to the integrated data. Automatic cell type annotation was executed with the help of SingleR[71] based on the mouse immune bulk expression data (ImmGenData), NK cells were manually confirmed by high expression of NK1.1 (Klrb1c) and NKp46 (Ncr1) and lacked expression of CD3 (Cd3e). Four sub-clusters of NK cells were obtained by re-applying UMAP analysis and clustering (with resolution set to 0.4) from the NK cells subset extracted from the integrated data. Gene set enrichment analysis was carried out by escape[72] package using AUCell score[73] to evaluate the activity of hallmark pathways and natural killer cells related GO biological process terms described in MSigDB[74, 75]. Regulon activity analysis was conducted with the help of the pySCENIC package according to the pipeline described before[73]. Differentially expressed genes were identified by FindMarkers in the Seurat package using the Wilcoxon Rank Sum test, genes with adjusted p value less than 0.05 and average foldchanges larger or less than 1 were considered significant. Cell-cell communication analysis was applied on each sample separately, single cell level expression profiles with gene symbols converted to human were fed through the liana[76] package that integrated results from different cell-cell communication detection methods and cell interaction resources. All single-cell plots including dimension plot, feature plot, gene expression boxplot, and violin plot were conducted by Scpub[77] package in R, and statistical tests for scRNA-seq data were carried out on R.

### Cytokine array

At various time points after *Salmonella* YB1 injection, mouse blood was collected to get serum. The cytokine level in murine serum was determined using a commercial cytokine array kit (Proteome Profiler Mouse XL Cytokine Array, R&D systems) according to the manufacturer’s instructions.

### Cytokine neutralization

To deplete cytokines *in vivo*, antibodies were used to neutralize IL-12 and IL-18. On day 0, BALB/c mice were administered *Salmonella* YB1. *In vivo* depletion of IL-12 was initiated one day before *Salmonella* YB1 injection through *i.p.* injection of 1mg of anti-IL-12 antibody (Cat.no BE0233; clone R2-9A5; Bioxcell), which was followed by additional *i.p.* injection on days 0 (1.2mg), 1 (0.3mg), 2 (0.3mg), 4 (0.3mg), 6 (0.2mg), 8 (0.2mg), 10 (0.2mg) and 12 (0.2mg). Rat IgG2b (Cat.no BE0090; clone LTF-2; Bioxcell) serves as a control. Similarly, *in vivo* depletion of IL-18 was initiated one day before *Salmonella* YB1 injection through *i.p.* injection of 500µg of anti-IL-18 antibody (Cat. No BE0237; clone YIGIF74-1G7; Bioxcell), which was followed by additional *i.p.* injection of anti-IL-18 antibody on days 0 (1.2mg), 1 (0.3mg), 2 (0.3mg), 4 (0.3mg), 6 (0.2mg), 8 (0.2mg), and 10 (0.2mg). Rat IgG2a (Cat.no BE0089; clone 2A3; Bioxcell) isotype control serves as a control. On day 30, 1.2×10^5^ 4T1 cells were injected into each mouse to develop lung metastasis.

### Cytokine treatment

On day 0, each BALB/c mouse received an intravenous injection of 10µg recombinant murine IL-12 (Cat.no 577008; Biolegend). IL-18 treatment was initiated through intraperitoneal injection of 2µg recombinant murine IL-18 (Cat.no 767006; Biolegend) on day 0, followed by additional intraperitoneal injections of recombinant murine IL-18 on days 1 (1µg), 2 (1µg), 3 (0.5µg), 4 (0.5µg), 5 (0.3µg), and 6 (0.3µg). IL-15 treatment was achieved through intraperitoneal injection of 100ng recombinant murine IL-15 (Cat.no 566302; Biolegend) on day 1 and day 2. 1.2×10^5^ 4T1 cells were injected into each mouse to develop lung metastasis on day 13 or day 33.

### PD-1 blockade

To assess the anti-metastatic ability of PD-1 blockade therapy, *in vivo* PD-1 blockade was initiated 2 days before 4T1 cancer cell inoculation (day 7) through *i.p.* injection of 250µg of anti-PD-1 antibody (Cat. BE0146; Bioxcell), followed by additional *i.p.* injection of anti-PD-1 antibody on days 7 (250µg), 9 (250µg), 12 (250µg), 15 (250µg), and 18 (200µg). To evaluate the impact of PD-1 blockade on early survival of 4T1-Luci cancer cells in the lungs, *in vivo* PD-1 blockade was initiated 3 days prior to inoculation of 4T1-Luci cancer cells via *i.p.* injection of 250µg of anti-PD-1 antibody at 4 hours after *Salmonella* YB1 treatment, followed by additional *i.p.* injection of 250µg of anti-PD-1 antibody 2 days later. Rat IgG2a (Cat.no BE0089; Bioxcell) served as an isotype control for PD-1 blockade.

### TIGIT blockade

To assess the anti-metastatic ability of TIGIT blockade therapy, *in vivo* TIGIT blockade was initiated 2 days before 4T1 cancer cell inoculation (day 0) through *i.p.* injection of 200µg of anti-TIGIT-1 antibody (Cat. BE0274; Bioxcell), followed by additional *i.p.* injection of anti-TIGIT antibody on days 0 (200µg), 2 (200µg), 5 (200µg), 8 (200µg), and 11 (200µg). Mouse IgG1 (Cat.no BE0083; Bioxcell) served as an isotype control for the TIGIT blockade.

### Statistical analysis

Subjects were allocated to groups using random assignment to minimize bias. To reduce potential bias, investigators and subjects were blinded during the conduct and analysis of the study. We referred to published work in similar field of study to determine appropriate group size for all experiments. Data were analyzed using GraphPad Prism version 9.0.2. The analyses are indicated in corresponding legends. All *P*-values are indicated in corresponding figures.

## Acknowledgments

We thank members of the Huang lab for helpful discussions and the Centre for PanorOmic Sciences at LKS Faculty of Medicine, the University of Hong Kong, for providing support on flow cytometry analysis and cell sorting. We thank Prof. Liwei LU (Department of Pathology, HKU) for sharing the CD45.1 transgenic mice (Pep Boy mice) with us. We thank Dr. Shachuan FENG and Miss. Jingchun MA for their critical suggestions on the manuscript writing. We thank Dr. Bin YU for genetically engineering *Salmonella* YB1. Icon for mouse from Figures 1a, 1i, 1l, 2f, 3i, 6f and Figures S2c, S3a, S3b, S6a, S7a, S8d, S9a, S9b, S9d was created by Dr. Li RONG. The research was supported by the National Key Research and Development Program of China (2018YFA0902701), J.D.H. thanks the L & T Charitable Foundation, the Program for Guangdong Introducing Innovative and Entrepreneurial Teams (2019BT02Y198), and Shenzhen Key Laboratory for Cancer Metastasis and Personalized Therapy (ZDSYS20210623091811035) for their support. The funding bodies did not contribute to the design of the study, or collection, analysis, and interpretation of the data.

## Data Availability Statement

The datasets we generated (ATAC-seq, CUT&RUN-seq, Bulk RNA-seq, and scRNA-seq) will be deposited at GEO(NCBI) and publicly available from the date of publication. All data and code supporting the findings of this study are available within the article and its supplementary information files. Any additional information required to reanalyze the data reported in this paper is available from the corresponding authors upon reasonable request.

## Author contributions

J.D.H. and L.R. conceived and designed the study; L.R., W.J., R.L., Y.W., H.S., X.W., S.H., H.G., X.L., W.L., Y.H., S.K., and X.Z. performed experiments; J.H. did all bioinformatics analysis and relevant writing work; N.Z., Z.Z., and Q.L. provided critical comments on the project; L.R. and J.D.H. analyzed the data and wrote the manuscript; N.Z. revised the manuscript; All authors read, commented on, and approved this manuscript. L.R., J.H., W.J., R.L., and Y.W. have been designated as co– first authors.

## Reference

1. Demaria, O., et al., Harnessing innate immunity in cancer therapy. Nature, 2019. 574(7776): p. 45–56.

2. Netea, M.G., et al., Defining trained immunity and its role in health and disease. Nat Rev Immunol, 2020. 20(6): p. 375–388.

3. Kaufmann, E., et al., BCG Educates Hematopoietic Stem Cells to Generate Protective Innate Immunity against Tuberculosis. Cell, 2018. 172(1-2): p. 176–190 e19.

4. Kleinnijenhuis, J., et al., Bacille Calmette-Guerin induces NOD2-dependent nonspecific protection from reinfection via epigenetic reprogramming of monocytes. Proc Natl Acad Sci U S A, 2012. 109(43): p. 17537–42.

5. Giamarellos-Bourboulis, E.J., et al., Activate: Randomized Clinical Trial of BCG Vaccination against Infection in the Elderly. Cell, 2020. 183(2): p. 315–323 e9.

6. de Laval, B., et al., C/EBPbeta-Dependent Epigenetic Memory Induces Trained Immunity in Hematopoietic Stem Cells. Cell Stem Cell, 2023. 30(1): p. 112.

7. Geller, A.E., et al., The induction of peripheral trained immunity in the pancreas incites anti-tumor activity to control pancreatic cancer progression. Nat Commun, 2022. 13(1): p. 759.

8. Singh, A.K., et al., Re-engineered BCG overexpressing cyclic di-AMP augments trained immunity and exhibits improved efficacy against bladder cancer. Nat Commun, 2022. 13(1): p. 878.

9. Wang, T., et al., Influenza-trained mucosal-resident alveolar macrophages confer long-term antitumor immunity in the lungs. Nat Immunol, 2023. 24(3): p. 423–438.

10. Broquet, A., et al., Sepsis-trained macrophages promote antitumoral tissue-resident T cells. Nat Immunol, 2024. 25(5): p. 802–819.

11. Artis, D. and H. Spits, The biology of innate lymphoid cells. Nature, 2015. 517(7534): p. 293-301.

12. Zhang, Y., et al., In vivo kinetics of human natural killer cells: the effects of ageing and acute and chronic viral infection. Immunology, 2007. 121(2): p. 258–65.

13. Chiossone, L., et al., Natural killer cells and other innate lymphoid cells in cancer. Nat Rev Immunol, 2018. 18(11): p. 671–688.

14. Cooper, M.A., et al., Cytokine-induced memory-like natural killer cells. Proc Natl Acad Sci U S A, 2009. 106(6): p. 1915–9.

15. Romee, R., et al., Cytokine activation induces human memory-like NK cells. Blood, 2012. 120(24): p. 4751–60.

16. Yu, B., et al., Explicit hypoxia targeting with tumor suppression by creating an “obligate” anaerobic Salmonella Typhimurium strain. Sci Rep, 2012. 2: p. 436.

17. Li, C.X., et al., ’Obligate’ anaerobic Salmonella strain YB1 suppresses liver tumor growth and metastasis in nude mice. Oncol Lett, 2017. 13(1): p. 177–183.

18. Lin, Q., et al., IFN-gamma-dependent NK cell activation is essential to metastasis suppression by engineered Salmonella. Nat Commun, 2021. 12(1): p. 2537.

19. Iwai, Y., S. Terawaki, and T. Honjo, PD-1 blockade inhibits hematogenous spread of poorly immunogenic tumor cells by enhanced recruitment of effector T cells. Int Immunol, 2005. 17(2): p. 133–44.

20. Zhang, Q., et al., Blockade of the checkpoint receptor TIGIT prevents NK cell exhaustion and elicits potent anti-tumor immunity. Nat Immunol, 2018. 19(7): p. 723–732.

21. Mombaerts, P., et al., RAG-1-deficient mice have no mature B and T lymphocytes. Cell, 1992. 68(5): p. 869–77.

22. Shultz, L.D., et al., Human lymphoid and myeloid cell development in NOD/LtSz-scid IL2R gamma null mice engrafted with mobilized human hemopoietic stem cells. J Immunol, 2005. 174(10): p. 6477–89.

23. Pantelouris, E.M., Absence of thymus in a mouse mutant. Nature, 1968. 217(5126): p. 370–1.

24. Pelleitier, M. and S. Montplaisir, The nude mouse: a model of deficient T-cell function. Methods Achiev Exp Pathol, 1975. 7: p. 149–66.

25. Hsu, J., et al., Contribution of NK cells to immunotherapy mediated by PD-1/PD-L1 blockade. J Clin Invest, 2018. 128(10): p. 4654–4668.

26. Lau, C.M., G.M. Wiedemann, and J.C. Sun, Epigenetic regulation of natural killer cell memory. Immunol Rev, 2022. 305(1): p. 90–110.

27. Cardoso, H.J., M.I. Figueira, and S. Socorro, The stem cell factor (SCF)/c-KIT signalling in testis and prostate cancer. J Cell Commun Signal, 2017. 11(4): p. 297–307.

28. Lee, S.H., M.F. Fragoso, and C.A. Biron, Cutting edge: a novel mechanism bridging innate and adaptive immunity: IL-12 induction of CD25 to form high-affinity IL-2 receptors on NK cells. J Immunol, 2012. 189(6): p. 2712–6.

29. Ni, J., et al., Sustained effector function of IL-12/15/18-preactivated NK cells against established tumors. J Exp Med, 2012. 209(13): p. 2351–65.

30. Stein, N., et al., IFNG-AS1 Enhances Interferon Gamma Production in Human Natural Killer Cells. iScience, 2019. 11: p. 466–473.

31. Ruckert, T., et al., Clonal expansion and epigenetic inheritance of long-lasting NK cell memory. Nat Immunol, 2022. 23(11): p. 1551–1563.

32. Beaulieu, A.M., et al., The transcription factor Zbtb32 controls the proliferative burst of virus-specific natural killer cells responding to infection. Nat Immunol, 2014. 15(6): p. 546–53.

33. Crinier, A., et al., High-Dimensional Single-Cell Analysis Identifies Organ-Specific Signatures and Conserved NK Cell Subsets in Humans and Mice. Immunity, 2018. 49(5): p. 971–986 e5.

34. Dean, I., et al., Rapid functional impairment of natural killer cells following tumor entry limits anti-tumor immunity. Nat Commun, 2024. 15(1): p. 683.

35. Reggiani, F., et al., BET inhibitors drive Natural Killer activation in non-small cell lung cancer via BRD4 and SMAD3. Nat Commun, 2024. 15(1): p. 2567.

36. Judge, S.J., W.J. Murphy, and R.J. Canter, Characterizing the Dysfunctional NK Cell: Assessing the Clinical Relevance of Exhaustion, Anergy, and Senescence. Front Cell Infect Microbiol, 2020. 10: p. 49.

37. Adams, N.M., et al., Transcription Factor IRF8 Orchestrates the Adaptive Natural Killer Cell Response. Immunity, 2018. 48(6): p. 1172–1182 e6.

38. Madera, S., et al., Type I IFN promotes NK cell expansion during viral infection by protecting NK cells against fratricide. J Exp Med, 2016. 213(2): p. 225–33.

39. Wang, K.S., J. Ritz, and D.A. Frank, IL-2 induces STAT4 activation in primary NK cells and NK cell lines, but not in T cells. J Immunol, 1999. 162(1): p. 299–304.

40. Wu, X., et al., Application of PD-1 Blockade in Cancer Immunotherapy. Comput Struct Biotechnol J, 2019. 17: p. 661–674.

41. Worboys, J.D., et al., TIGIT can inhibit T cell activation via ligation-induced nanoclusters, independent of CD226 co-stimulation. Nat Commun, 2023. 14(1): p. 5016.

42. Ochando, J., et al., Trained immunity - basic concepts and contributions to immunopathology. Nat Rev Nephrol, 2023. 19(1): p. 23–37.

43. Cheng, S.C., et al., Broad defects in the energy metabolism of leukocytes underlie immunoparalysis in sepsis. Nat Immunol, 2016. 17(4): p. 406–13.

44. Roquilly, A., et al., Alveolar macrophages are epigenetically altered after inflammation, leading to long-term lung immunoparalysis. Nat Immunol, 2020. 21(6): p. 636–648.

45. Bauer, M., et al., Remembering Pathogen Dose: Long-Term Adaptation in Innate Immunity. Trends Immunol, 2018. 39(6): p. 438–445.

46. Morvan, M.G. and L.L. Lanier, NK cells and cancer: you can teach innate cells new tricks. Nat Rev Cancer, 2016. 16(1): p. 7–19.

47. Arase, H., et al., Direct recognition of cytomegalovirus by activating and inhibitory NK cell receptors. Science, 2002. 296(5571): p. 1323-6.

48. Nabekura, T., et al., Costimulatory molecule DNAM-1 is essential for optimal differentiation of memory natural killer cells during mouse cytomegalovirus infection. Immunity, 2014. 40(2): p. 225–34.

49. Sun, J.C., et al., Proinflammatory cytokine signaling required for the generation of natural killer cell memory. J Exp Med, 2012. 209(5): p. 947–54.

50. Madera, S. and J.C. Sun, Cutting edge: stage-specific requirement of IL-18 for antiviral NK cell expansion. J Immunol, 2015. 194(4): p. 1408–12.

51. Nabekura, T., J.P. Girard, and L.L. Lanier, IL-33 receptor ST2 amplifies the expansion of NK cells and enhances host defense during mouse cytomegalovirus infection. J Immunol, 2015. 194(12): p. 5948–52.

52. Ni, J., et al., Adoptively transferred natural killer cells maintain long-term antitumor activity by epigenetic imprinting and CD4(+) T cell help. Oncoimmunology, 2016. 5(9): p. e1219009.

53. Anderson, R.L., et al., A framework for the development of effective anti-metastatic agents. Nat Rev Clin Oncol, 2019. 16(3): p. 185–204.

54. Patro, R., et al., Salmon provides fast and bias-aware quantification of transcript expression. Nat Methods, 2017. 14(4): p. 417–419.

55. Love, M.I., W. Huber, and S. Anders, Moderated estimation of fold change and dispersion for RNA-seq data with DESeq2. Genome Biol, 2014. 15(12): p. 550.

56. Wu, T., et al., clusterProfiler 4.0: A universal enrichment tool for interpreting omics data. Innovation (Camb), 2021. 2(3): p. 100141.

57. Corces, M.R., et al., An improved ATAC-seq protocol reduces background and enables interrogation of frozen tissues. Nat Methods, 2017. 14(10): p. 959–962.

58. Kaya-Okur, H.S., et al., CUT&Tag for efficient epigenomic profiling of small samples and single cells. Nat Commun, 2019. 10(1): p. 1930.

59. Chen, S., et al., fastp: an ultra-fast all-in-one FASTQ preprocessor. Bioinformatics, 2018. 34(17): p. i884–i890.

60. Langmead, B. and S.L. Salzberg, Fast gapped-read alignment with Bowtie 2. Nat Methods, 2012. 9(4): p. 357–9.

61. Tarasov, A., et al., Sambamba: fast processing of NGS alignment formats. Bioinformatics, 2015. 31(12): p. 2032–4.

62. Ramirez, F., et al., deepTools2: a next generation web server for deep-sequencing data analysis. Nucleic Acids Res, 2016. 44(W1): p. W160–5.

63. Amemiya, H.M., A. Kundaje, and A.P. Boyle, The ENCODE Blacklist: Identification of Problematic Regions of the Genome. Sci Rep, 2019. 9(1): p. 9354.

64. Stark, R. and G. Brown, DiffBind: differential binding analysis of ChIP-Seq peak data. R package version, 2011. 100(4.3).

65. Fornes, O., et al., JASPAR 2020: update of the open-access database of transcription factor binding profiles. Nucleic Acids Res, 2020. 48(D1): p. D87–D92.

66. Schep, A., motifmatchr: fast motif matching in R. R package version 1.26. 0. 2024.

67. Heinz, S., et al., Simple combinations of lineage-determining transcription factors prime cis-regulatory elements required for macrophage and B cell identities. Mol Cell, 2010. 38(4): p. 576–89.

68. Li, D., et al., WashU Epigenome Browser update 2019. Nucleic Acids Res, 2019. 47(W1): p. W158–W165.

69. Zheng, G.X., et al., Massively parallel digital transcriptional profiling of single cells. Nat Commun, 2017. 8: p. 14049.

70. Hao, Y., et al., Dictionary learning for integrative, multimodal and scalable single-cell analysis. Nat Biotechnol, 2024. 42(2): p. 293–304.

71. Aran, D., et al., Reference-based analysis of lung single-cell sequencing reveals a transitional profibrotic macrophage. Nat Immunol, 2019. 20(2): p. 163–172.

72. Borcherding, N., et al., Mapping the immune environment in clear cell renal carcinoma by single-cell genomics. Commun Biol, 2021. 4(1): p. 122.

73. Aibar, S., et al., SCENIC: single-cell regulatory network inference and clustering. Nat Methods, 2017. 14(11): p. 1083–1086.

74. Subramanian, A., et al., Gene set enrichment analysis: a knowledge-based approach for interpreting genome-wide expression profiles. Proc Natl Acad Sci U S A, 2005. 102(43): p. 15545–50.

75. Castanza, A.S., et al., Extending support for mouse data in the Molecular Signatures Database (MSigDB). Nat Methods, 2023. 20(11): p. 1619–1620.

76. Dimitrov, D., et al., Comparison of methods and resources for cell-cell communication inference from single-cell RNA-Seq data. Nat Commun, 2022. 13(1): p. 3224.

77. Blanco-Carmona, E., Generating publication ready visualizations for Single Cell transcriptomics using SCpubr. bioRxiv, 2022: p. 2022.02.28.482303.

