## Supporting Information for "Therapeutic *Salmonella* induces long-term protective trained immunity in NK cells against cancer metastasis"

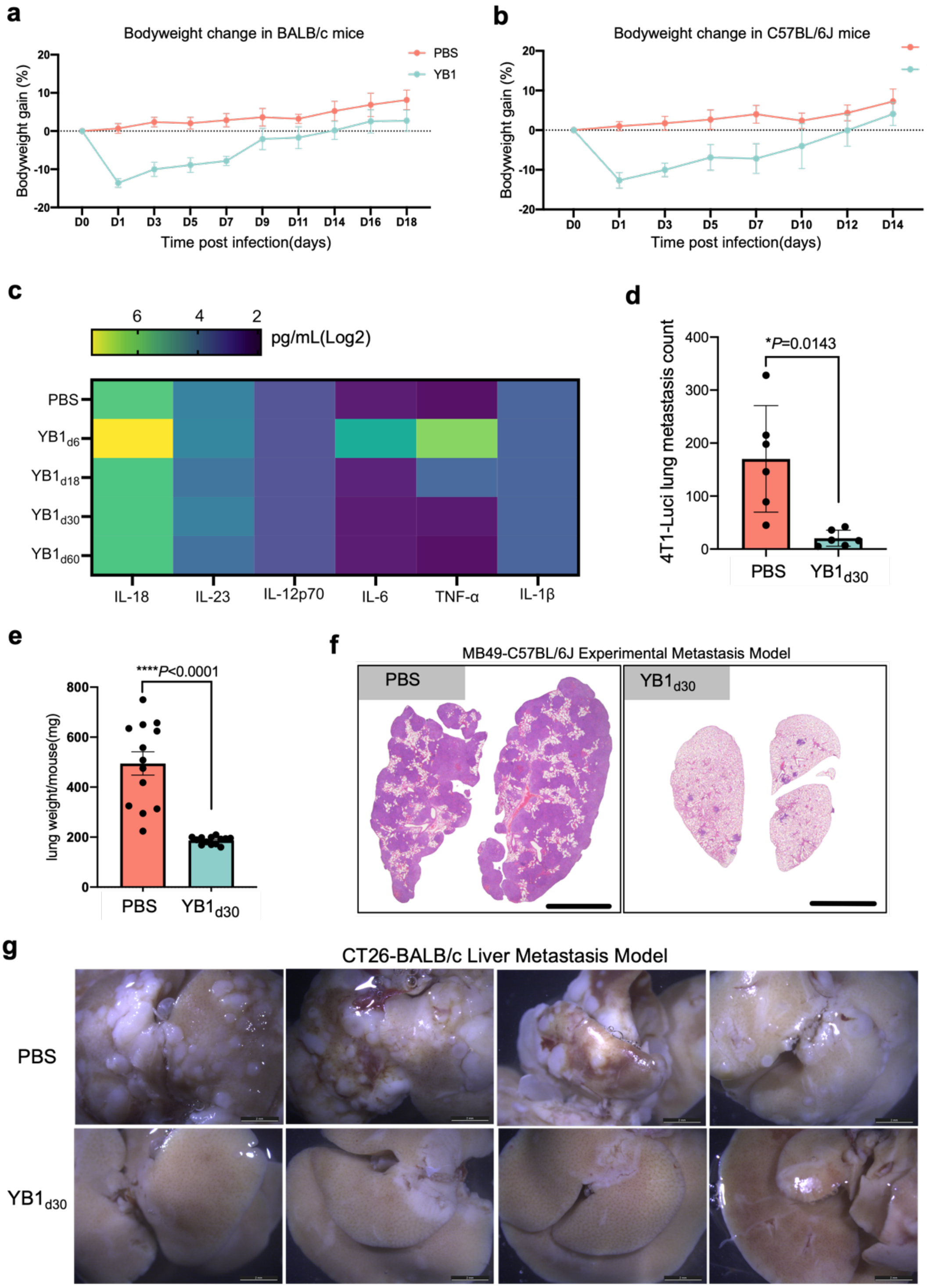


**Figure S1. Attenuated Salmonella treatment suppresses metastasis in multiple syngeneic mouse tumor models after the resolution of infection-induced inflammation (related to Figure 1).**

**a)** BALB/c mice were either treated with PBS (n=6) or 2x10$^7$ CFU of *Salmonella* YB1 (n=6), respectively. Daily changes of body weight were monitored for both groups.

**b)** C57BL/6J mice were either treated with PBS (n=10) or 2x$10^7$ CFU of *Salmonella* YB1 (n=13), respectively. Daily changes of body weight were monitored for both groups.

**c)** Quantitative analysis of serum cytokine levels in BALB/c mice (n=4) treated with PBS or YB1 at various time points post-infection.

**d)** Quantification of 4T1-Luci cell lung metastases (two-tailed unpaired t-tests, n=6 per group, displayed one representative experiment of two independent experiments, presented as mean values ± SD) 14 days after the luciferase live imaging, as depicted in **Figure 1e, f**.

**e-f)** Quantification of lung weight (unpaired two-tailed t-test, combined results of 2 independent experiments, n=13 per group, presented as mean values ± SEM) and representative whole lung H&E staining (scale bar, 3mm) in the same cohort of mice as depicted in **Figure 1g, h**.

**g)** Representative liver photos (scale bar, 2mm) showing CT26 liver metastatic nodules as depicted in **Figure 1i-k**.

All *P* values are shown in the relevant figures.

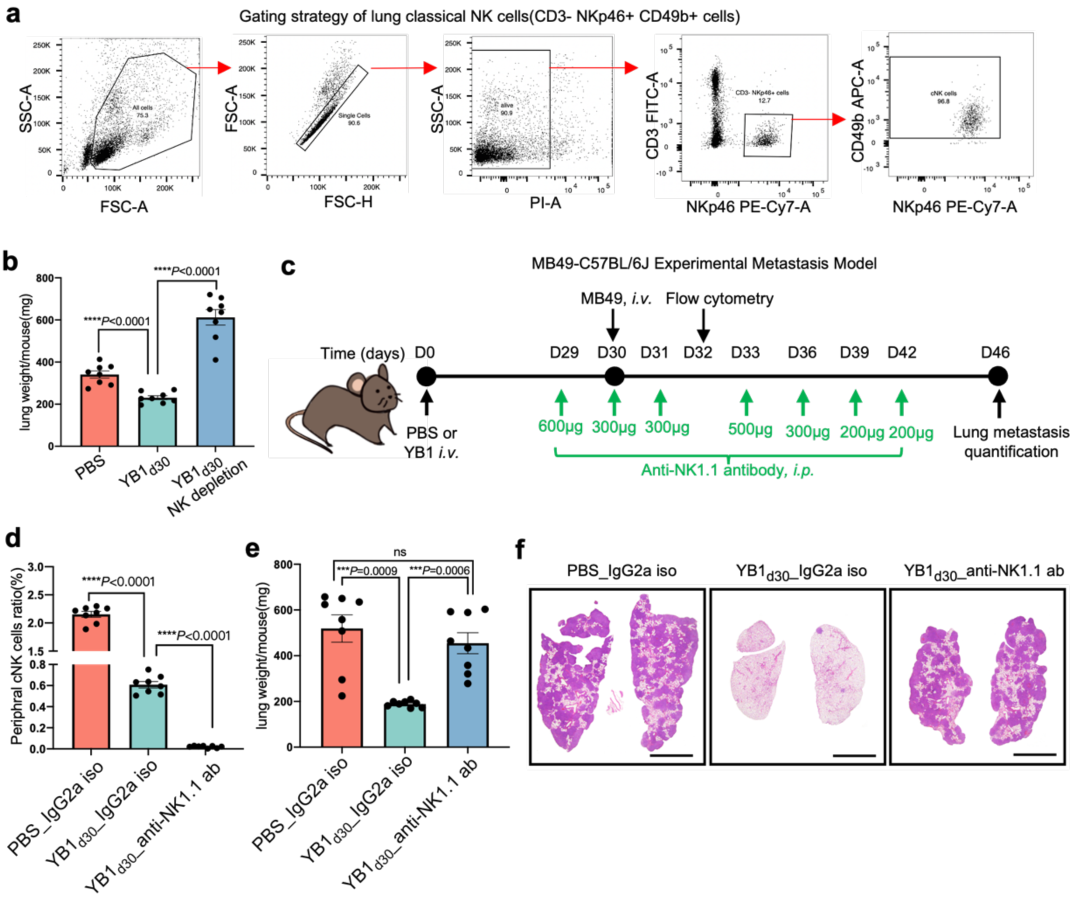


**Figure S2. NK cells mediate the long-lasting anti-metastatic immunity conferred by attenuated *Salmonella* (related to Figure 2)**

**a)** General gating strategy for flow analysis. Gates applied in every experiment: SSC-A/FSC-A gate was used to remove debris, FSC-A/FSC-H was used to gate singlets, and PI-positive gate was used to remove dead cells if not for intracellular staining. After this, cells were detected with fluorescent signals of relevant channels. For example, CD3- NKp46+ gate and further CD49b+ gate were used to define NK cells in BALB/c mice, while CD3- NK1.1+ gate and further CD49b+ gate were used to define NK cells in C57BL/6J mice.

**b)** Quantification of lung weight (n=8 per group) in the same cohort of BALB/c mice as depicted in **Figure 2f-i**.

**c)** A schematic representation of NK cell depletion schedule using Anti-NK1.1 antibody in C57BL/6J mice pretreated with YB1 30 days prior. MB49 cancer cells were *i.v.* injected into C57BL/6J mice 30 days post-treatment with either YB1 or PBS, and the NK cell depletion efficiency was measured by flow cytometry.

**d)** The peripheral ratio of NK cells in C57BL/6J mice was validated by flow cytometry (n=8 per group) 2 days post MB49 cancer cell inoculation, as depicted in **Figure S2c.**

**e-f)** Quantification of lung weight (n=8 per group) and representative whole lung H&E staining (scale bar, 3mm) in the same cohort of mice as depicted in **Figure S2c** and **Figure 2j-k**.

All quantified data are shown as combined results of 2 independent experiments and presented as mean values ± SEM unless indicated. All *P* values are yielded by two-tailed unpaired t-tests and are shown in the relevant figures.

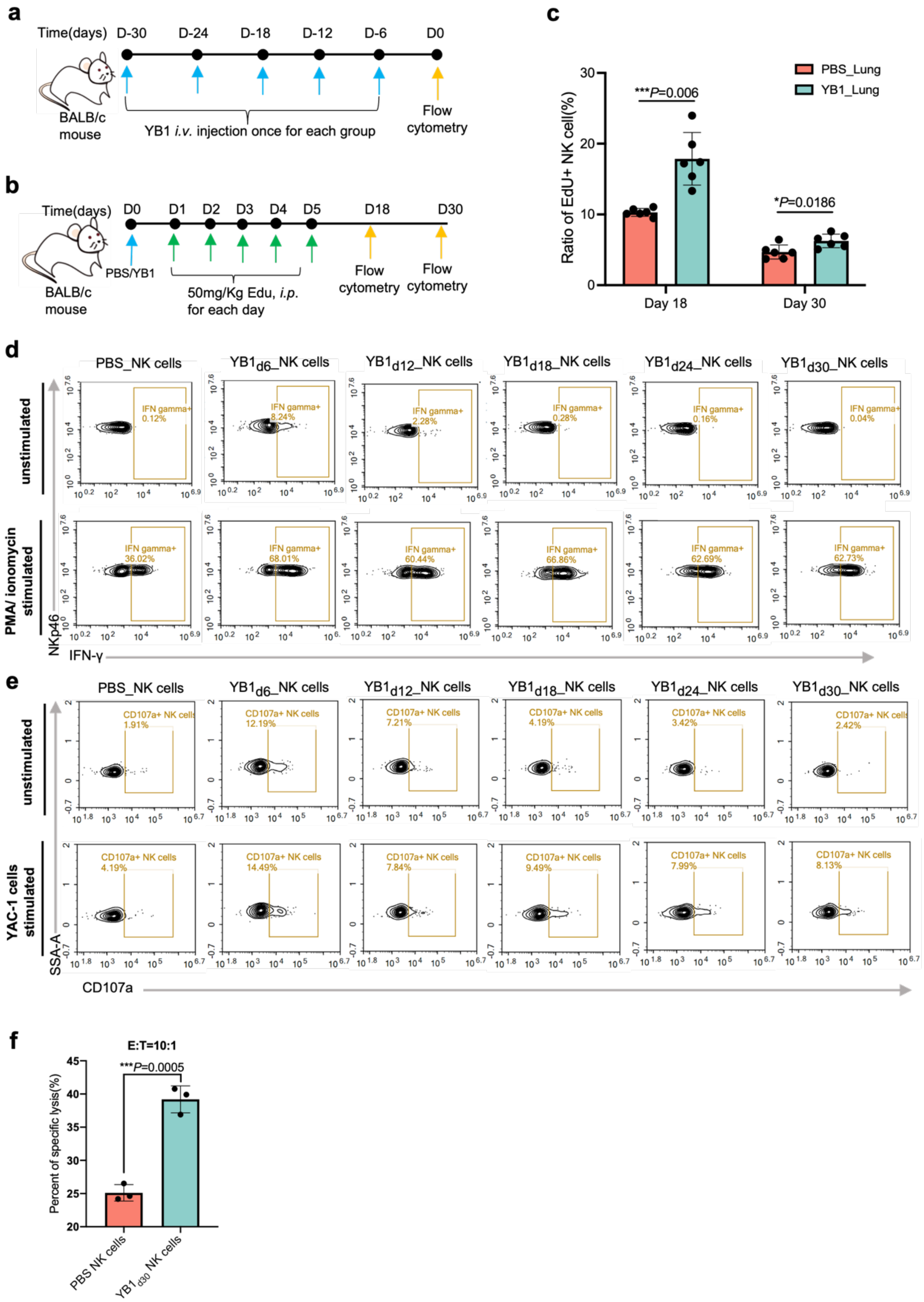


**Figure S3. stNK cells exhibit enhanced cytokine production and cytotoxicity upon secondary stimulation after return to homeostasis (related to Figure 3a-h)**

**a)** A schematic representation of flow cytometry analysis of peripheral and lung infiltrating immune cells, as shown in **Figure 3d-g**. BALB/c mice were pre-treated with YB1 at various time points before flow cytometry analysis.

**b)** A schematic representation of EdU incorporation post-treatment with either YB1 or PBS. Each mouse received a daily intraperitoneal injection of 50 mg/kg of EdU for 5 consecutive days to label proliferating NK cells, which was followed by assessing the proportion of EdU+ NK cells to total NK cells 18 or 30 days later. PBS serves as a control treatment for YB1 treatment.

**c)** The quantification (two-tailed unpaired t-tests, n =6 per group, Combined results of 2 independent experiments, data are presented as mean values ± SEM) of EdU+ NK cells to total NK cells in the lung across samples as depicted in **Figure S3b**.

**d)** Representative flow plots display IFN-γ and NKp46 expression in lung-infiltrating NK cells after *ex vivo* culture without or with PMA/Ionomycin stimulation for 5 hours. Quantification of the same sample batch can be found in **Figure 3f**.

**e)** Representative flow plots exhibit CD107a and SSA expression in lung-infiltrating NK cells after *ex vivo* culture without or with YAC-1 tumor cells for 5 hours. Quantification of the same sample batch is presented in **Figure 3g**.

**f)** Cytotoxicity of lung-infiltrating NK cells against YAC-1 target cells is shown at the indicated NK cells: YAC-1 cells (E: T, Effector: Target) ratio of 10:1 (two-tailed unpaired t-tests, n=3 biological replicates, displaying one representative experiment of two independent experiments, data are presented as mean values ± SD).

Lung-infiltrating immune cells (**Figure S3d, e**) or NK cells (**Figure S3f)** were isolated from BALB/c mice pre-treated with *Salmonella* YB1 at different time points, with PBS-treated mice serving as controls. All *P* values are yielded by two-tailed unpaired t-tests and are shown in the relevant figures.

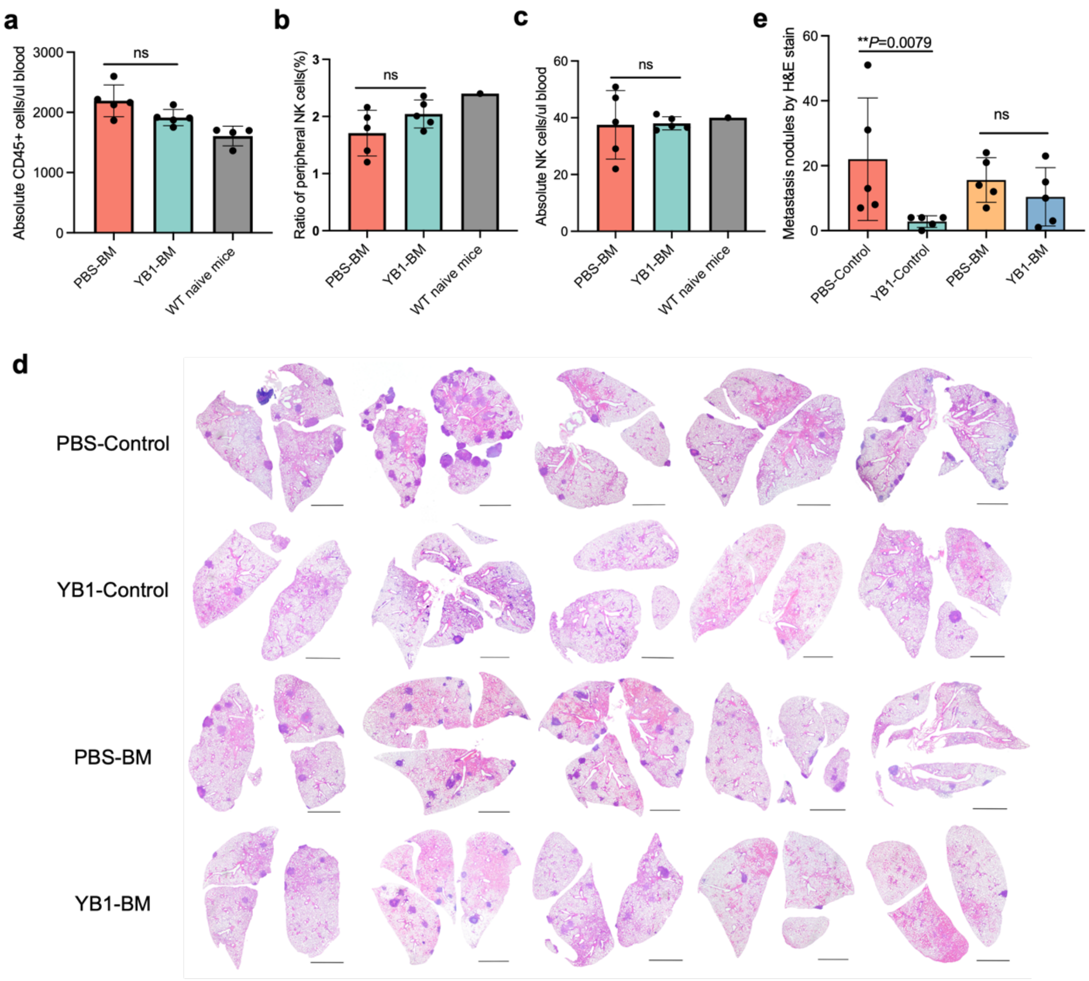


**Figure S4. Attenuated *Salmonella* treatment doesn't induce central anti-metastatic trained immunity (related to Figure 3i-k)**

Bone marrow cells were extracted from C57BL/6N mice 23 days after PBS or YB1 treatment and transplanted into naïve, irradiated, syngeneic CD45.1 mice. Flow cytometry analysis was performed on day 58 (5 weeks after bone marrow transplantation) to assess bone marrow reconstitution efficiency. On day 61, recipient mice and control C57BL/6N mice were injected intravenously with MB49 bladder cancer cells to induce lung metastasis, as depicted in **Figure 3i**.

**a)** Measurement of total CD45+ cell concentration (cells/µL of blood) following a 5-week bone marrow reconstitution period. The gray bar indicates wild-type naïve mice control.

**b)** The peripheral ratio of donor-derived NK cells following a 5-week bone marrow reconstitution period. The gray bar indicates a wild-type naïve mouse control.

**c)** Measurement of donor-derived NK cell concentration (cells/µL of blood) following a 5-week bone marrow reconstitution period. The gray bar indicates a wild-type naïve mouse control.

**d-e)** Representative H&E staining of whole lung (scale bar, 2mm) from different groups and its quantification of tumor nodules.

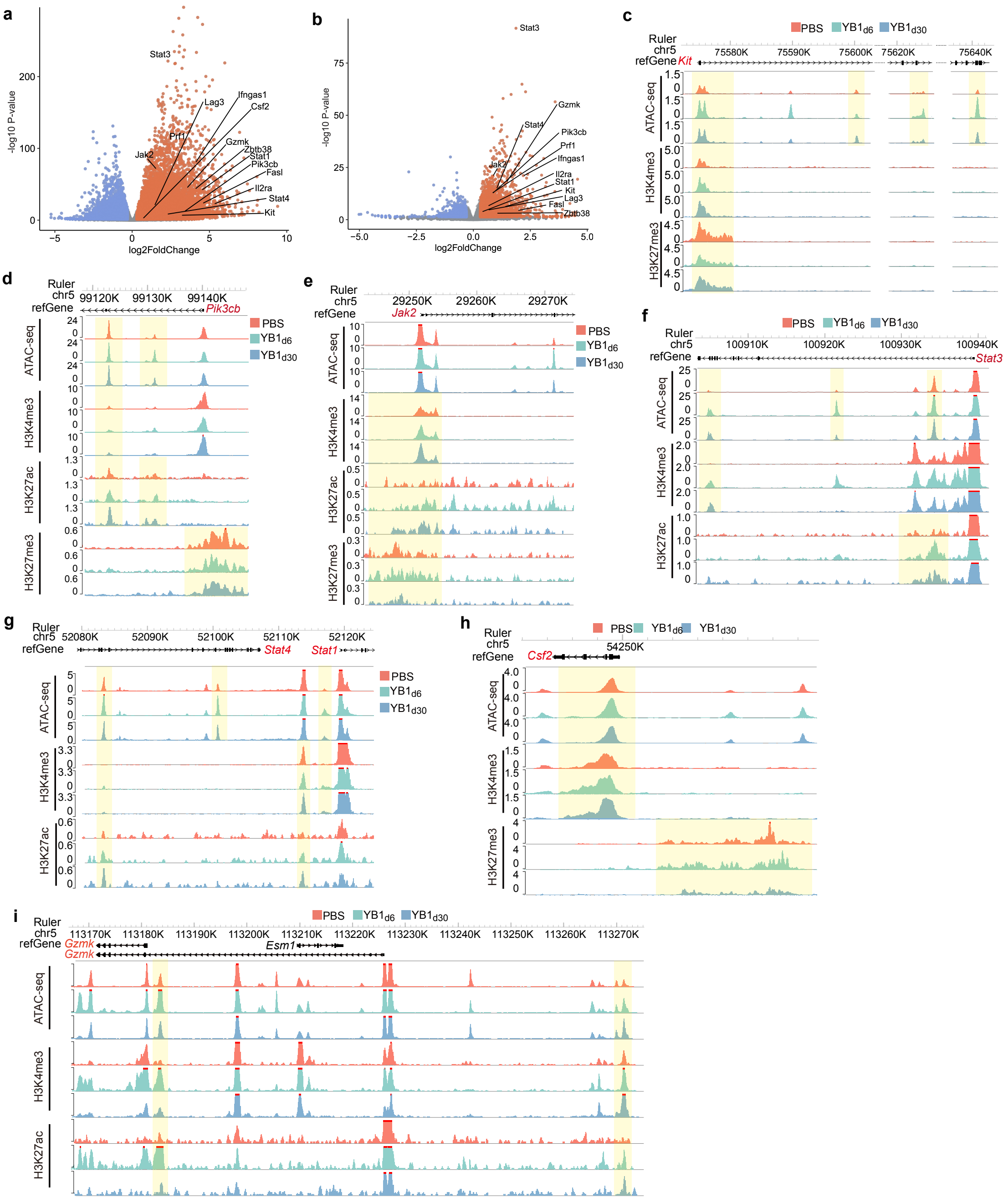

**Figure S5. Therapeutic Salmonella induces long-lasting epigenetic changes in NK cells (related to Figure 4)**

**a-b)** Volcano plots display differential chromatin accessibility regions in stNK cells 6 days (**a**) or 30 days (**b**) post-infection compared to control NK cells. Labeled genes of interest are shown.

**c-i**) Genome browser tracks of interested gene regions, such as *Kit* (**c**), *Pik3cb* (**d**), *Jak2* (**e**), *Stat3* (**f**), *Stat1*(**g**), *Csf2* (**h**) and *Gzmk* (**i**) in control NK cells and stNK cells (6 days and 30 days post-infection) are presented. Gene regions exhibiting epigenetic changes suggestive of increased transcriptional accessibility are indicated by shadow boxes.

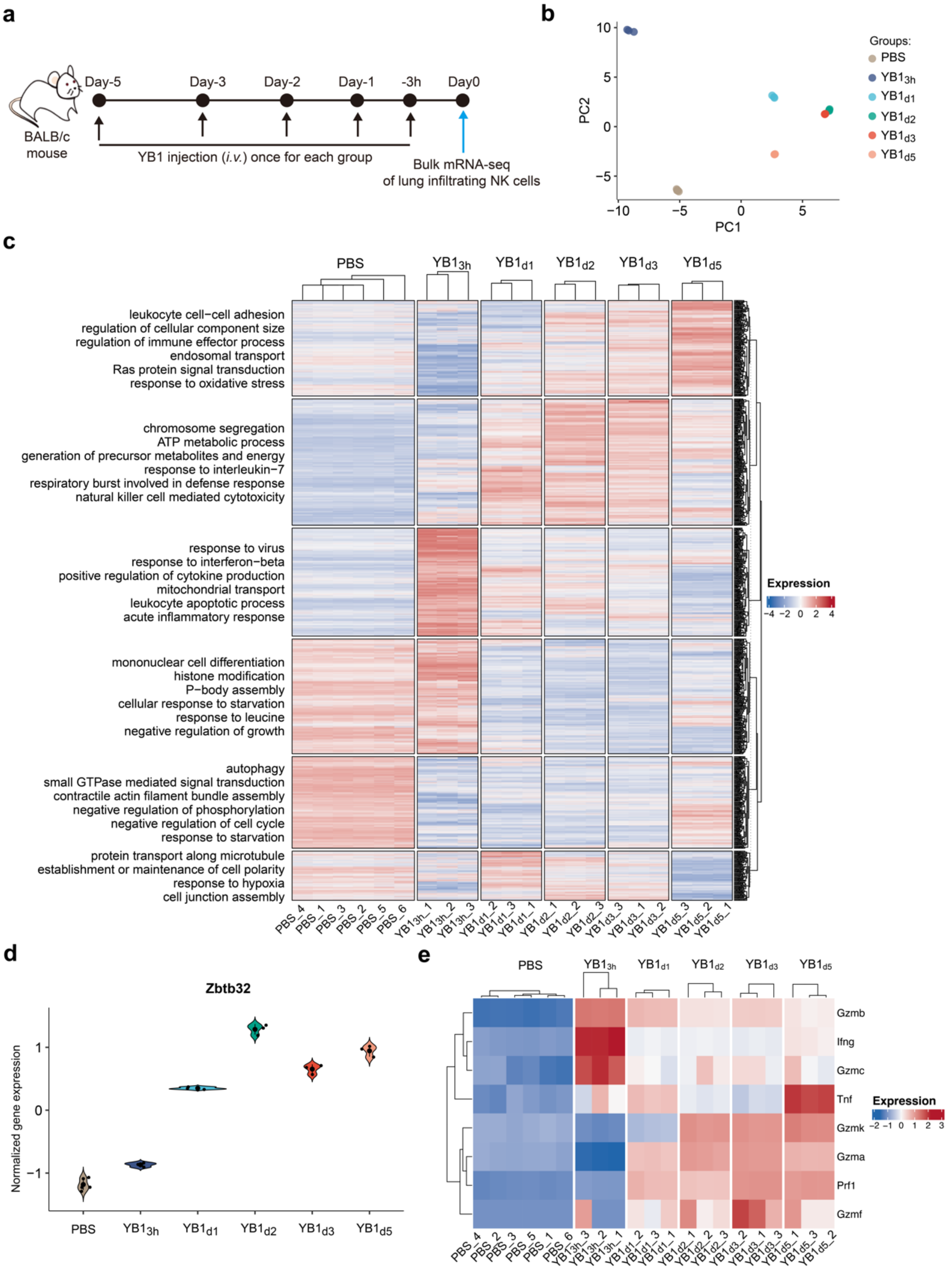


**Figure S6. Bulk RNA-seq analysis of lung-infiltrating NK cells during the acute phase of *Salmonella* infection (related to Figure 5)**

**a)** A schematic representation of the experimental design for the bulk RNA-seq analysis is shown. Lung-infiltrating NK cells were isolated from BALB/c mice pre-treated with YB1 at various time points prior. PBS serves as a control treatment.

**b)** PCA of bulk RNA-seq data is displayed.

**c)** Hierarchical clustering of DEGs was performed based on normalized expressions. Only genes with significant upregulation compared to the PBS group (adjusted p-value $\leq 0.05$ and log2 fold changes $\geq 1$) at any time points after YB1 treatment were included. The top enriched gene ontology terms for each gene cluster are indicated on the left.

**d)** The normalized expression of gene *Zbtb32* is presented.

**e)** Normalized expression of selected immune effector genes is shown.

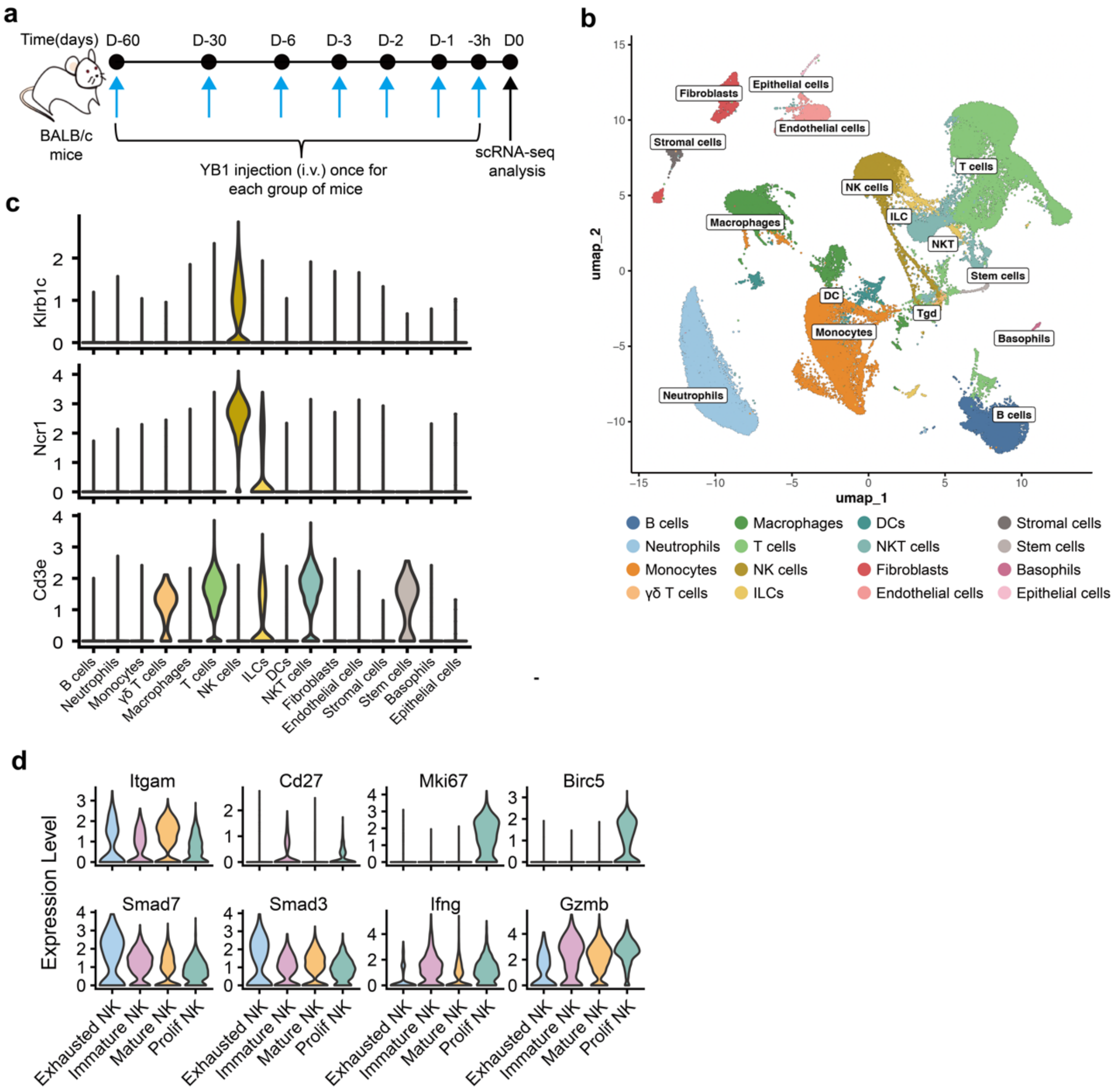


**Figure S7. scRNA-seq analysis of lung-infiltrating immune cells (related to Figure 5)**

**a)** A schematic representation of the experimental design for scRNA-seq is shown. Lung-infiltrating immune cells were isolated from YB1-treated mice at various time points post-YB1 infection. PBS serves as a control treatment.

**b)** UMAP dimensionality reduction plot of all cells is displayed.

**c)** Expression profiles of markers used for NK cell population annotation are presented.
**d)** Expression profiles of marker genes used for NK cell subtype annotation are shown.

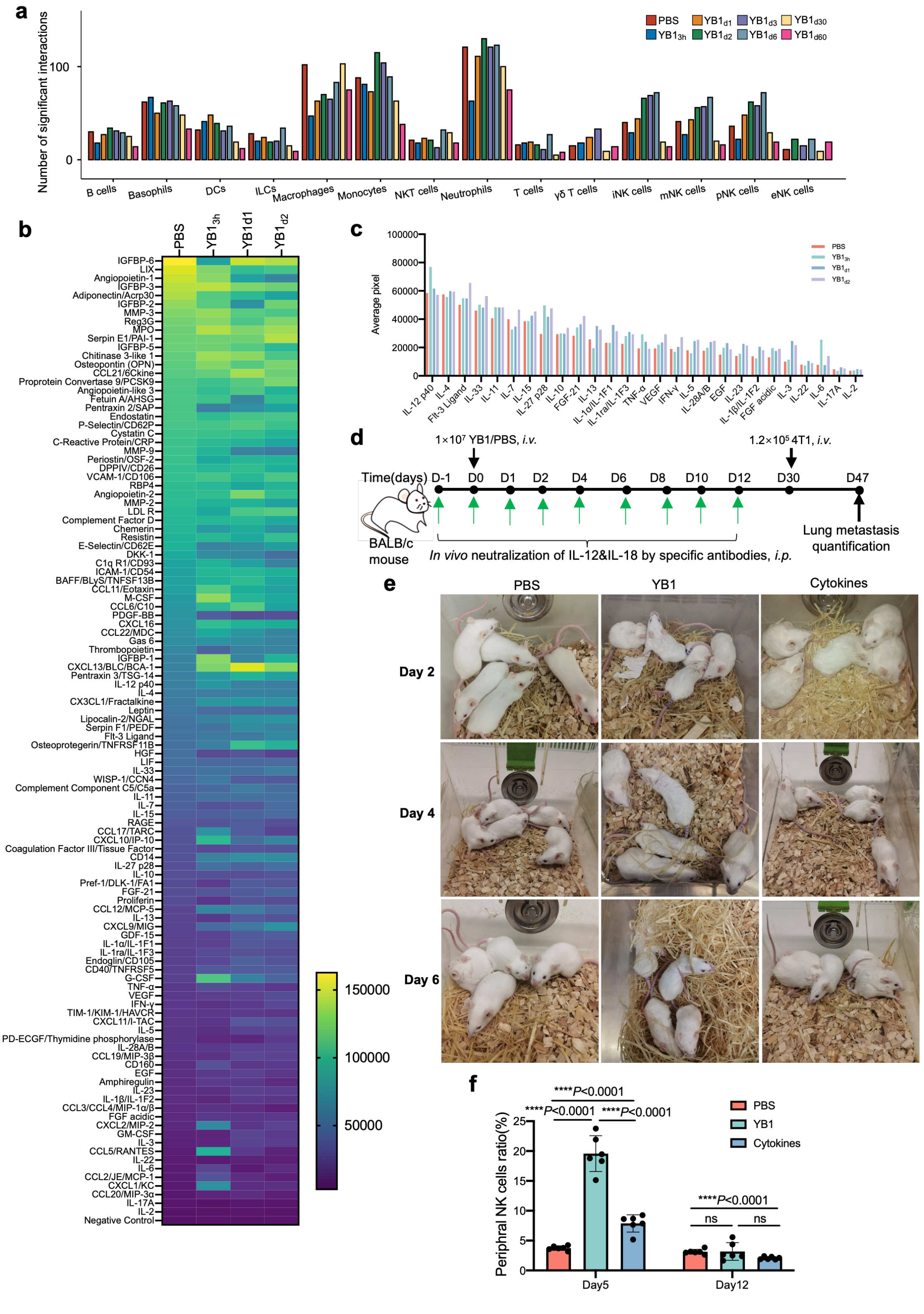


**Figure S8. IL-12 and IL-18 are necessary for the development of stNK cells (related to Figure 6)**

**a)** The number of interactions of different immune cells to NK cells. pNK: Prolif NK, iNK: Immature NK, mNK: Mature NK, eNK: Exhausted NK.

**b)** The cytokine array heatmap displays the dynamics of all tested cytokines and chemokines in the serum of mice treated with PBS or YB1 at various time points post-infection.

**c)** The bar graph of selected cytokine array results as depicted in **Figure 6c**.

**d)** The experimental design for *in vivo* neutralization of IL-12 and IL-18 induced by YB1 treatment is schematically represented. PBS- or YB1-treated mice served as controls.

**e)** Photos of mice from different groups on days 2, 4, and 6.

**f)** The percentage (two-tailed unpaired t-tests, n=6 per group, combined results of 2 independent experiments, represented as mean values ± SEM) of peripheral NK cells in differently treated mice: PBS, YB1, or cytokines.

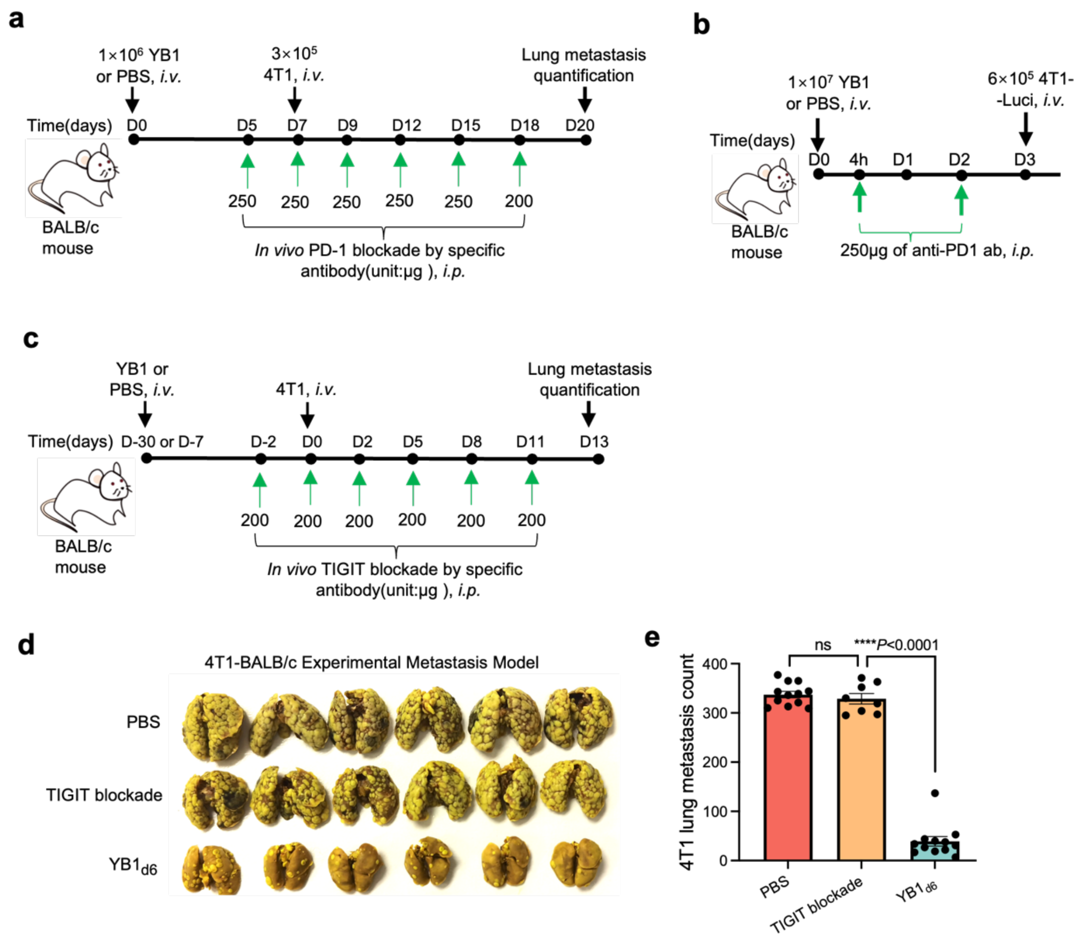


**Figure S9. Attenuated *Salmonella* outplays immune checkpoint blockade therapies in suppressing metastasis (related to Figure 7)**

**a)** A schematic representation of the experimental design comparing the anti-metastatic effects of PD-1 blockade and YB1 treatment is shown. The results are presented in **Figure 7a, b**.

**b)** A schematic representation of the experimental design comparing the inhibitory effects of PD-1 blockade and YB1 treatment on early cancer cell survival in the lungs is displayed. The results are shown in **Figure 7c, d**.

**c)** A schematic representation of the experimental design comparing the anti-metastatic effects of TIGIT blockade and YB1 treatment is presented. The results are presented in **Figure 7e, f** as well as in **Figure S9d, e**.

**d-e)** Images of Bouin-fixed lungs (**d**) and the quantification (**e**) of 4T1 lung metastases (n=12 for PBS and YB1$_{d6}$ groups, n=8 for TIGIT blockade group, combined results of 2 independent experiments, presented as mean values ± SEM) for the mice receiving different treatments, as depicted in **Figure S9c**.

All *P* values are yielded by two-tailed unpaired t-tests and are shown in the relevant figures.

**Table S1 Antibodies for flow cytometry analysis**

| Antigen | Fluorescent conjugate | Species reactivity | Vendor | Catalog Number | Clone |
|---|---|---|---|---|---|
| CD3 | FITC | mouse | eBioscience | 11-0032-82 | 17A2 |
| CD3 | Alexa Fluor 700 | mouse | BioLegend | 100216 | 17A2 |
| CD49b | APC | mouse | BioLegend | 108910 | DX5 |
| NKp46 | PE-Cy7 | mouse | eBioscience | 25-3351-82 | 29A1.4 |
| NK1.1 | PE-Cy7 | mouse | BioLegend | 156514 | S17016D |
| NKp46 | PE | mouse | eBioscience | 12-3351-80 | 29A1.4 |
| CD11b | eFluor 450 | mouse | eBioscience | 48-0112-80 | M1/70 |
| CD27 | Super Bright 600 | mouse | Thermo Fisher Scientific | 63-0271-80 | LG.7F9 |
| CD11c | Alexa Fluor 700 | mouse | eBioscience | 56-0114-80 | N418 |
| IFN-γ | APC | mouse | eBioscience | 17-7311-81 | XMG1.2 |
| CD3 | Brilliant Violet 785 | mouse | BioLegend | 100232 | 17A2 |
| CD107a | V450 | mouse | BD Biosciences | 560648 | 1D4B |
| CD11b | Pacific blue | mouse & human | BioLegend | 101224 | M1/70 |
| TIGIT | APC | Mouse | BioLegend | 142105 | 1G9 |
| PD-1 | PerCP-eFluor 710 | Mouse | Thermo Scientific | 46-9985-80 | J43 |
| CD45 | APC | mouse | BioLegend | 103112 | 30-F11 |
| CD11c | FITC | mouse | BioLegend | 117306 | N418 |
| F4/80 | Alexa Fluor 700 | mouse | Thermo Scientific | 56-4801-80 | BM8 |
| CD45.2 | Pacific blue | mouse | BioLegend | 109820 | 104 |
| CD45.1 | PE-Cy7 | mouse | thermo fisher scientific | 25-0453-81 | A20 |
| NK1.1 | Brilliant Violet 711 | mouse | Biolegend | 108745 | PK136 |
